# Aberrant activation of Wnt/β-Catenin signaling pathway drives the expression of poor prognosis-associated microRNAs in adrenocortical cancer with a major impact on miR-139-5p and its host gene PDE2A

**DOI:** 10.1101/2023.02.10.527992

**Authors:** Justine Cristante, Soha Reda El Sayed, Josiane Denis, Bruno Ragazzon, Constanze Hantel, Olivier Chabre, Laurent Guyon, Nadia Cherradi

## Abstract

Adrenocortical carcinoma (ACC) is a rare malignancy with dismal prognosis. Deregulated microRNA (miRNA) expression has been implicated in ACC aggressiveness. Nevertheless, the mechanisms underlying such deregulations remain unknown. Aberrant Wnt/β-Catenin signaling has been reported in about 40% of ACC and is associated with poor outcome. Here, we investigated the link between constitutive activation of Wnt/β-Catenin pathway and miRNA expression alterations in ACC. Inducible shRNA-mediated gene silencing of β-Catenin (β-Cat) was performed in ACC cells expressing constitutively active β-Catenin. The miRnome of ACC cells was analyzed using RNA-Sequencing. Selected miRNAs and mRNAs were validated using quantitative PCR and functional experiments with an emphasis on miR-139-5p, its host gene phosphodiesterase 2A (PDE2A) and its target gene N-Myc Downstream-Regulated Gene 4 (NDRG4). Prognostic values of Wnt/β-Catenin pathway components or mutational status and their correlations with miRNA/mRNA expressions were determined in COMETE-ENSAT and TCGA cohorts. We carried out the first miRnome analysis in β-Catenin-deficient (β-Cat^-^) ACC cells. Twelve upregulated miRNAs and 42 downregulated miRNAs among which miR-139-5p and miRNAs of the 14q32 locus were identified in β-Cat^-^ cells. Downregulation of selected poor prognosis-associated miRNAs was confirmed using RT-qPCR. Remarkably, the expression of the intronic miR-139-5p was decreased by 90% in β-Cat^-^ cells with a concomitant repression of its host gene PDE2A and upregulation of its target gene NDRG4. In ACC patients, miR-139-5p levels were highly correlated with the levels of PDE2A and anti-correlated with those of NDRG4. MiR-139-5p and PDE2A expressions were higher in patients with mutations in components of Wnt/β-Catenin signaling pathway or high expression of LEF1, with LEF1 proving a better predictor of prognosis than Wnt/β-Catenin signaling pathway mutational status. Our findings indicate that in addition to inducing protein-coding genes in ACC, constitutively active Wnt/β-Catenin signaling upregulates the expression of a subset of miRNAs involved in tumour aggressiveness and poor clinical outcome.

## Introduction

Adrenocortical carcinoma (ACC) is a rare cancer of the adrenal cortex, with an incidence of 0.7-2 cases/millions of persons/year in adults and an incidence peak between 50 and 60 years.^1^ 50 to 70% of patients present hormonal disorders, mainly hypercortisolism (Cushing’s syndrome) or androgen excess.^2^ However, incidental discovery of ACC on abdominal computed tomography (incidentalomas) is increasing, and is now estimated around 15%.^3^ Patients are classified according to clinical stages, with stages I and II representing localized tumour, while stages III and IV represent advanced pathology with lymph node or systemic metastasis. ACC prognosis is very poor, with a 5-year survival between 12 and 40% and a median survival below 3 years.^4^ Overall survival is significantly different depending on the stage with a 5-year survival of 65-82% for stage I, 58-64% for stage II, 44-55% for stage III and 7-18% for stage IV.^5^ 50 to 71% of ACC patients are diagnosed at late stages.^6^ To date, treatment is based on complete surgical resection for localized ACC followed by mitotane adjuvant therapy.^3^ In advanced or recurrent ACC, the therapeutic arsenal is limited to repeated surgery, local therapeutic measures and chemotherapy, but it remains with poor efficacy. Targeted therapies and immunotherapy for ACC have been used in recent clinical trials with limited benefits.^7^

Two major studies deciphered the genomic landscape of ACC using the French COMETE/European ENSAT cohort^8^ and the American TCGA cohort.^9^ Transcriptome unsupervised clustering identified two subgroups of tumors C1A and C1B with very poor and much better outcomes, respectively.^8^ Among the major molecular alterations found in ACC, somatic mutations in *CTNNB1* gene, encoding β-Catenin, and in *ZNRF3* gene, encoding an E3 ubiquitin-protein ligase that acts as a repressor of Wnt/β-Catenin pathway,^10, 11^ were the most frequent and mutually exclusive (16% and 21%, respectively).^8, 9^ This results in Wnt/β-Catenin signaling being constitutively active in more than 35% of ACC. The canonical Wnt/β-Catenin signaling is a highly conserved pathway, which plays a major role in cell growth, development and oncogenesis.^12^ In ACC, Wnt/β-Catenin pathway is mostly activated in aggressive tumors and is associated with poor prognosis.^8, 13, 14^ Diffuse abnormal cytoplasmic and nuclear β-Catenin localization was found both in patient tumors and in the human ACC cell line H295R.^15^ Genetic analyses revealed that the most commonly detected mutation is the point mutation S45P (exon 3) affecting the consensus GSK3β phosphorylation site,^15^ thus leading to β-Catenin stabilization and constitutive activation of T-cell factor/lymphoid enhancer-binding factor (TCF/LEF)-dependent transcription of its target genes including *AXIN2, MYC* and *LEF1*. β-Catenin silencing in H295R cells resulted in a decreased proliferation *in vitro*, abrogated tumor growth *in vivo*^*16*^ and led to the down-regulation of the transcription factor Slug,^17^ which is associated with a more aggressive phenotype of ACC.^18^

Besides alterations of the Wnt/β-Catenin signaling pathway, massive deregulations of microRNA (miRNA) expression have been identified in ACC.^8, 9, 19, 20^ MiRNAs are small non-coding RNA of about 22 nucleotides-long,^21^ which repress their target gene expression at the post-transcriptional level, either by inducing mRNA cleavage, destabilization or inhibition of translation.^22^ MiRNAs are deregulated in cancer and are involved in carcinogenesis and drug resistance. They also emerged as promising biomarkers for cancer diagnosis and prognosis^23^ and potential therapeutic tools.^24^ Commonly overexpressed miRNAs in ACC include miR-483, miR-139-5p, miR-503, miR-184 and miR-210, while miR-195 and miR-335 were found underexpressed in ACC compared to adrenocortical adenoma (ACA) or normal tissue.^19^ In addition, some miRNA clusters have been reported as particularly deregulated between poor and good prognosis ACC, such as the 14q32 or the Xq27 clusters.^8, 25^ We have previously shown that miR-139-5p and miR-483-5p promote ACC aggressiveness, at least by respectively downregulating the N-myc Downstream-Regulated Gene member 4 (*NDRG4*) and member 2 (*NDRG2*).^26^

MiRNAs have been widely described as direct or indirect regulators of Wnt/β-Catenin signaling pathway in several cancers such as clear cell renal cell carcinoma,^27^ colorectal cancer,^28^ hepatocellular carcinoma,^29^ and breast cancer.^30^ By contrast, only very few studies have examined modulations of miRNA expression in response to active Wnt/β-Catenin pathway. Herein, we report on the impact of aberrant activation of Wnt/β-Catenin signaling on the whole miRnome in ACC. We show that silencing of β-Catenin leads to significant downregulation of several poor prognosis-associated oncomiRs in ACC including miR-139-5p and miRNAs located in the 14q32 cluster. We identified miR-139-5p and its host gene Phosphodiesterase 2A (*PDE2A*) as major targets of the Wnt signaling pathway components β-Catenin and Lymphoid enhancer-binding factor 1 (LEF1) transcription factor. The expression of miR-139-5p in ACC cells as well as in ACC patients was strongly correlated with the expression of *PDE2A*, suggesting a common regulatory transcription unit. Furthermore, we provide evidence that LEF1 is a significant prognostic factor for ACC. Besides, its high expression is better associated with miR-139-5p and *PDE2A* upregulation than the mutational status of *CTNNB1*/*β-Catenin* or *ZNRF3*.

## Materials and methods

### Cell culture and transfection

The adrenocortical carcinoma cell lines H295R (ATCC, #CRL-2128) and MUC-1 MUC were subjected to Cell Line Sterility, IMPACT Mouse FELASA and MKPV testing and was authenticated by CellCheck human 9 Test (9 Marker STR Profile and Inter-species Contamination Test) (IDEXX Analytics, Kornwestheim, Germany). H295R cells stably expressing a doxycycline inducible β-Catenin shRNA (pTer1 shRNA vector, shβ7 cells) or pTer1 control vector (pTer1 cells)^16^ were authenticated by CellCheck human 16 Plus Test (9 human STR marker profile + 7 additional Human STR marker, Inter-species contamination Test and STAT-mycoplasma testing) (IDEXX Analytics). They were grown in Dulbecco’s Modified Eagle Medium:F-12 (DMEM/F-12 Glutamax, Gibco Life Technologies, Carlsbad, USA, #31331093) supplemented with 1% (v/v) Insulin, Transferrin, Selenium Premix (Corning, NY, USA, #354352), 5% (v/v) Nu Serum (Corning, #355100) and antibiotics (100 U/mL penicillin, 100 µg/mL streptomycin and 30 µg/mL gentamicin, Life Technologies). Petri dishes were pre-coated with Rat Tail Collagen Type 1α (Corning, #354236, 1:50 dilution). Stable *CTNNB1/β-Catenin* silencing was achieved through shRNA induction in the presence of doxycycline (0.5µg/mL) (H295R β-Cat^-^ cells), while β-Catenin expression was maintained in the absence of doxycycline (H295R β-Cat^+^ cells).

Kinetics of gene expression were conducted for 10 days with molecular analyses at day 0 (D0), day 3 (D3), day 5 (D5), day 7 (D7) and day 10 (D10). Briefly, shβ7 and pTer1 cells were seeded in petri dishes of 100mm of diameter, at a density of 2 × 10^6^ cells, incubated the following day (D0) in complete medium with or without Doxycycline and lysed at the appropriate day.

For transient silencing of *CTNNB1/β-Catenin*, H295R cells were seeded in 6-wells plate at a density of 0.3×10^6^ cells/well and transfected the day after with a *CTNNB1* siRNA (Thermofisher, Carlsbad, USA, #s436) or a control siRNA (Thermofisher, #AM4611), at a final concentration of 10nM, using Lipofectamine RNAiMAX (Invitrogen Thermofisher, Carlsbad, USA, #13778150) according to the manufacturer’s recommendations.

MUC1 cells have been generated previously from an ACC neck metastasis of a male patient.^31^ They were grown in Advanced DMEM-F12 (Gibco Thermofisher, Carlsbad, USA, #12634-010) with 1% Penicillin-Streptomycin 10UI/mL (Gibco Thermofisher, #15140-122) and 10% Fetal Bovine Serum (Biosera, Nuaille, France, #FB-1001/100). For comparison of H295R and MUC1 cell lines, cells were seeded in 6-well plates at a density of 0.7 × 10^6^ and 0.25 × 10^6^ cells/well, respectively. The following day (D0), medium was changed and cells were grown up for 72h before lysis.

For miR-139-5p rescue experiments, H295R shβ7 cells and unmodified H295R cells were seeded in 6-well plates at a density of 0.35 × 10^6^ cells/well and 0.3 × 10^6^ cells/ wells respectively. The following day, shβ7 cells were incubated in complete medium with or without Doxycycline. H295R cells were transfected with a *CTNNB1* siRNA or a control siRNA as described above. The same day, both shβ7 cells and H295R cells were transfected with control miRNA mimics or hsa-miR-139-5p mimics (Thermofisher, #MC11749) at a final concentration of 10nM. The transfection of miRNA mimics together with *CTNNB1* or control siRNAs was repeated at D6 for both cell lines. Cells were lysed at D10.

### Small RNA sequencing

Shβ7-expressing cells were grown for 10 days in a medium containing or not doxycyline. Total RNA was extracted using the miRVana™ Paris™ kit (Thermofisher, #AM1556) and depleted from ribosomal rRNA using Ribo-zero Removal kit (Illumina). RNA quality and quantity were assessed by SS RNA kit on Fragment Analyzer (Agilent) and QuantiFluor RNA System (Promega) respectively, according to the manufacturers’ instructions. Small RNA library preparation was performed using TruSeq Small RNA library preparation kit (Illumina) according to the manufacturer’s instructions. The quality and quantity of the obtained library were assessed using HS DNA chip on Bioanalyzer 2100 (Agilent) and QuantiFluor DNA System (Promega), respectively. Small RNA sequencing was performed on the Illumina NextSeq platform of the Centre National de Recherche sur le Genome (Evry, France), using MID-output flow cell to generate 51bp long reads.

### RNA sequencing data analysis

FASTQ data were processed as follows using Galaxy v20.01. The data were filtered using FastQC v0.72^32^ and Trim Galore! v0.4.3.1,^33^ aligned against the miRBase v22 sequence list^34^ using Bowtie1 v1.2.0,^35^ and then the read counts were normalized using DESeq2 v2.11.40.^36^ We then selected miRNAs according to three criteria using R version 4.0.3.^37^ MiRNAs were considered as expressed in sufficient quantity in cells when the number of read per millions (RPM) was above 10 (log2 RPM > 5.55) in at least one experimental condition (β-Cat^+^ or β-Cat^-^). MiRNAs were considered as differentially expressed between β-Cat^+^ and β-Cat^-^ cells when the log fold change (LFC) was > 2 or < -2. For each miRNA, the p-value for prognosis was calculated from the ENSAT Cohort data,^10^ with the log-rank test applied on two equal subpopulation separated with the tumour expression of the miRNA, using the R library “survival” version 3.2.^38^ We selected miRNAs with p-value below 0.05.

### Reverse Transcription - Quantitative PCR

Total RNA was extracted from shβ7 and pTer1 cells using miRVana™ Paris™ kit according to the manufacturer’s instructions. For miRNA quantification, 10 ng of total RNA were reverse-transcribed as previously described^26^ using TaqMan™ MicroRNA Reverse Transcription Kit (#4366596, Applied Biosystems Thermofisher) and miRNA-specific stem-loop primers (Applied Biosystems Thermofisher). Real-time quantitative PCR (qPCR) was performed using 4.5µL of 5-fold diluted RT product and 5.5µL of reaction mix (composed of 5µL of TaqMan Universal Master Mix (Applied Biosystems, Life Technologies, Carlsbad, USA, #4440047) with 0.5µL of the appropriate assay for each miRNA (miR-139-5p, miR-483-5p, miR-483-3p, miR-513a-5p, miR-377-5p, miR-377-3p, miR-1185-2-3p, miR-541-3p, miR-320b, miR-2110 and miR-330-3p, (Supplementary Table 1) as previously reported.^26^ RNU48 was used as an endogenous control for normalization. All experiments were performed in duplicate.

Messenger RNA expression was assessed with 150 to 500 ng of total RNA using iScript cDNA Synthesis Kit (Bio-Rad, Hercules, USA, #1708891) according to manufacturer’s instructions. Quantitative PCR was performed using 4.5µl of 1:30 diluted RT product and 5.5 µl of reaction mix (composed of 5 µl of Sso Advanced Universal SYBR Green Supermix (Bio-Rad, #1725275) and 0.5 µl of forward and reverse PCR primers mixed together at a final primer concentration of 0.5 µM) to generate a PCR volume of 10 µl. All primers were purchased from Sigma-Aldrich and are reported in Supplementary Table 2. Real-time PCR was carried out at 95°C for 30s, followed by 40 cycles at 95°C for 5s and 60°C for 15s. Gene expression was normalized to RPL13A and HPRT as housekeeping genes. Normalized expression was calculated using the comparative Ct method and fold changes were derived from the 2^-ΔΔCt^ values for each gene.

### Western blot

Proteins were extracted from shβ7 and pTer1 cells using Cell Disruption Buffer (miRVana™ Paris™ kit, Thermofisher) and were quantified with microBCA protein Assay kit (Thermofisher, #23235). Protein extracts were dissolved in 5X buffer (60 mM Tris-HCl, pH 6.8, 10% glycerol, 2 % SDS, 5 % β-mercapto-ethanol, 0.01% bromophenol blue), heated at 100°C for 5 min and loaded onto 4-20% mini-protean TGX precast protein gels. Electrophoresis was carried out at 80 V for 20-30 min and then at 120 V for 1 hour in a 10X Tris/glycine/SDS running buffer. SDS-PAGE resolved proteins were transferred onto a nitrocellulose membrane using the Trans-Blot Turbo™ transfer system (Bio-Rad) according to the manufacturer’s instructions. The membrane was then incubated for 1 hour at room temperature in a blocking solution of Tris-buffer saline (TBS) containing 0.1% Tween 20 and 5% non-fat dry milk.

The blots were probed overnight at 4°C with monoclonal mouse anti-β-Catenin (2 µg/mL in Bovine Serum Albumin 3%, R&D Systems, Minneapolis, USA, #MAB1329), monoclonal rabbit anti-NDRG4 (1:1000, Cell Signaling Technology, Danvers, USA, #9039), monoclonal rabbit anti-Axin2 (1:1000, #2151, Cell Signaling), polyclonal rabbit anti-PDE2A (1:1000, Sigma-Aldrich, #ABN1486) or monoclonal mouse anti-tubulin IgG (1:2500, Sigma-Aldrich, #T6199) in TBS/Tween. The membrane was then washed with the same buffer (3 × 10 min) and incubated for 1 hour either with Horseradish-Peroxydase (HRP) goat anti-rabbit IgG (1:3000, Jackson Immunoresearch, Cambridge, UK, #111-035-144) or with HRP Goat anti-mouse IgG (1:5000, Jackson Immunoresearch, #115-035-062). The membrane was washed as previously mentioned and the antigen-antibody complex was revealed by Clarity Enhanced Chemiluminescence Blotting Substrate (Bio-Rad, #1705060) using the iBright™ FL100 Imaging System (ThermoFisher). Signal intensity was quantified using the iBright Analysis Software (versions 4.0.1 and 5.0.0, Thermofisher). Protein levels were normalized to tubulin. Data are presented as the fold-change in protein expression relative to controls.

### Measurement of Wnt/β-Catenin signaling pathway activity (TOPflash luciferase reporter assay)

H295R shβ7 cells and pTer1 cells were seeded in triplicates in a 12-well plate (200 000 cells/well) and were incubated in the presence or in the absence of Doxycycline the day after. 24 hours later, 250 ng of pTOPflash (wt-TCF) or pFOPflash (mutated-TCF binding site, negative control) reporter plasmids (TOPflash luciferase reporter assay, Merck Millipore, Darmstadt, Germany) were transfected with Lipofectamine 2000 (Invitrogen, Life Technologies) in the presence of 150 ng of pRL-TK-Renilla luciferase plasmid (Promega, Charbonnières-les-Bains, France). After 48h incubation with Doxycycline, Firefly and Renilla luciferase activities were measured using the Twinlite dual luciferase reporter gene assay system (Perkin Elmer, Wellesley, MA, USA) on an automated TECAN luminometer (TECAN Group Ltd. Zürich, Switzerland). The ratio of Firefly luciferase activity (Relative light units RLU)/Renilla luciferase activity (RLU) was expressed as a fold change relative to the controls. H295R cells were seeded in triplicates into 12 well-plates (300 000 cells/well) and transfected for 48h with 10nM antimiR-139-5p using Lipofectamine RNAimax. After 2 days, pTOPflash or pFOPflash and pRL-TK-Renilla plasmids were transfected using Lipofectamine 2000 as described above. Firefly and Renilla luciferase activities were measured 24h after plasmid transfection.

### COMETE-ENSAT and TCGA patient cohort analyses

Gene expression analysis of the two cohorts were performed with R (version 4.0.3).^37^ For the COMETE-ENSAT cohort, public data from Assie *et al*.^8^ were extracted from GSE49276, GSE49277 and the supplementary data. Final analysis was performed on the 44 ACC for which we had both miRNAs and mRNA data. Patients with mutations in *CTNNB1* gene, *ZNRF3* gene or mutation in either gene in addition to *TP53* were classified as mutated for the Wnt/β-Catenin signaling pathway. Patients with other mutations were classified as non-mutated for the Wnt/β-Catenin signaling pathway.

For the TCGA cohort, data were retrieved from the TCGA website https://portal.gdc.cancer.gov/projects/TCGA-ACC. Expression data for miRNAs and mRNAs were available for 80 and 79 patients, respectively. The analysis was performed for the 79 patients for whom we had both miRNA and mRNA data. The “transcript per million” column (“tmp_unstranded” column) was used for mRNA analysis. Both miRNA and mRNA data were log2-transformed.

As the distribution of gene expression in the tumour samples of patients was bimodal, in particular for LEF1, we used the expectation-maximization (EM) algorithm to fit a Gaussian mixture using the R library “mixtools” version 1.2. To set the threshold between high and low LEF1 expression, the LEF1 expression with the same probability density for both groups was used, leading to a threshold of 5.2 for the COMET-ENSAT cohort.

### Statistical analyses

Statistical analyses were performed with R (version 4.0.3)^37^. Data normality was assessed using Shapiro-Wilk test with Benjamini-Hochberg correction for multiple tests. T-tests were used for mean comparison while Wilcoxon test was used if the data were not normal. Results are expressed as mean ± SEM. Statistical significance is indicated with * for p ≤ 0.05, ** for p ≤ 0.01 and *** for p ≤ 0.001. For qPCR, paired t-tests were performed on raw data of qPCR expression extracted from the Bio-Rad CFX Software, to compare control samples and doxycycline-treated samples. As our results indicated a stabilized *CTNNB1/β-Catenin* silencing at D2-D3, we assessed the effect of β-Catenin repression for long-term global kinetics, regardless of the precise day. Thus, we performed the statistics on pooled data from D5, D7 and D10. For each day, the β-Cat^-^ sample was paired to the β-Cat^+^ control of the same day. Bar plots represent the mean fold change in β-Cat^-^ samples for each day.

For western blot analyses, paired t-tests were used to compare the mean of doxycycline-treated samples with control samples, for each day. Gene expression fold-change in doxycycline-treated samples was normalized to that in control sample at each day.

## Results

### *β*-Catenin shRNA-mediated inactivation of Wnt signaling

We first assessed *β-Catenin* (*β-Cat*) silencing in H295R shβ7 cells upon shRNA induction by doxycycline. A marked reduction in β-Catenin mRNA (74 ± 11 %) was observed at day 3 (D3), reaching 91 ± 2 % at day 10 (D10) while a decrease of β-Catenin protein was detected at D3 (81 ± 3 %) and persisted until D10 (Figure 1a and 1b). No changes in β-Catenin expression occurred in the pTer1 control cells treated or not with doxycycline (Supplementary Figures S1a, 1b and 1e). As shown in Figure 1f, the activity of β-Catenin signaling pathway, assessed by the TOP-FOP Flash luciferase reporter assay, was also markedly decreased 48 h after β-Catenin silencing (96 ± 0.1 %), while no change was observed in pTer1 cells (Supplementary Figure S2).

**Figure 1.**
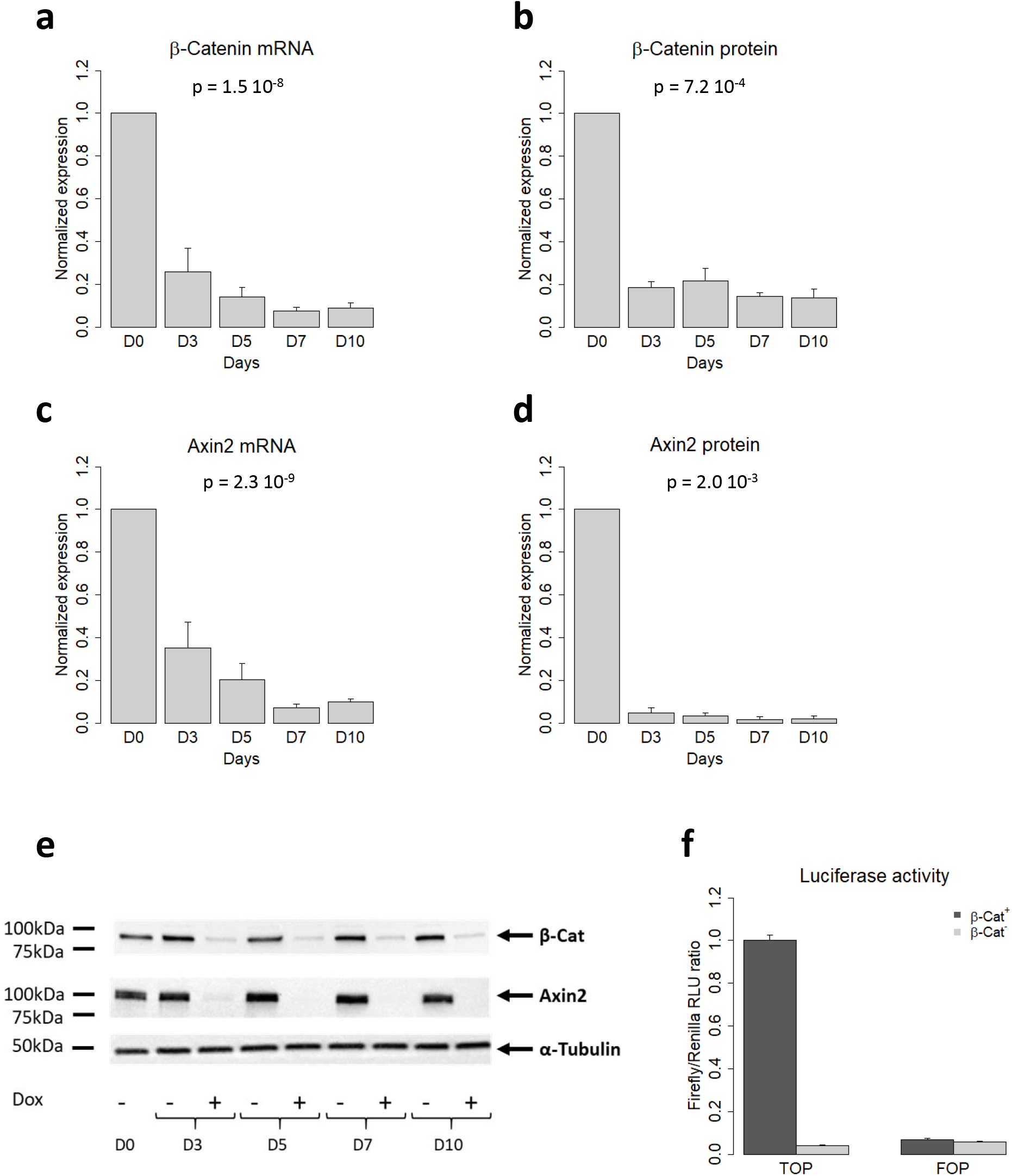
Doxycycline efficiently silences Wnt/β-Catenin signalling in H295R shβ7 cells. Cells were cultured without (β-Cat^+^) or with Doxycycline (β-Cat^-^) during 10 days. Cells were lysed at the day (D) indicated (D0, D3, D5, D7 and D10). (a) and (b) Expression levels of β-Catenin mRNA and protein, assessed by RT-qPCR and western blot, respectively, in β-Cat^-^ cells. (c) and (d) Expression levels of Axin2 mRNA and Axin2 protein, assessed by RT-qPCR and western blot, respectively, in β-Cat^-^ cells. (e) A representative western blot for β-Catenin and Axin2 protein levels in non-treated or doxycycline-treated cells. α-tubulin was used as a loading control. (f) Wnt/β-Catenin signaling TOP-flash reporter activity in β-Cat^+^ and β-Cat^-^ cells. The TOPflash reporter contains wild-type TCF/LEF binding sites while the FOPflash reporter contains the mutated TCF/LEF binding sites (negative control). Shown are the means ± S.E.M of 5 independent experiments performed in duplicate for qPCR analyses, 5 independent experiments for western blot analyses of β-Catenin and 4 independent experiments for Axin2.

The levels of AXIN2, a canonical target gene of β-Catenin, were also determined to validate the shβ7 cell model. As shown in Figure 1c, *AXIN2* mRNA level was decreased by 65 ± 12 % at D3 and 90 ± 2 % at D10. Axin2 protein expression was decreased by 95 ± 2 % at D3 and this decrease was maintained up to D10 (Figure 1d). No change in *AXIN2*/Axin2 occurred in the pTer1 control cells (Supplementary Figures S1c, 1d and 1e). Altogether, these results demonstrate that shRNA-mediated silencing of *β-Catenin* efficiently impairs Wnt signaling pathway activity.

### Inactivation of Wnt/β-Catenin signaling affects microRNA expression levels in adrenocortical carcinoma cells

To determine the impact of β-Catenin silencing on miRNA expression in H295R cells, we performed a miRNA profiling analysis using small RNA sequencing. The expression of 54 mature miRNAs was changed following β-Catenin repression (Figure 2a, Supplementary Table 3): 42 miRNAs were downregulated (77.8%) and 12 were upregulated in β-Cat^-^ cells compared to β-Cat^+^ cells. MiR-139-5p, that we have previously described as a marker of tumor aggressiveness,^25, 26^ was markedly repressed following silencing of β-Catenin, while miR-483-5p, a marker of malignancy, was not affected. Analysis of the genomic position of β-Catenin-regulated miRNAs using miRViz software^39^ revealed that many of them were localized in the same genomic region (Figure 2b), namely the 14q32 locus that we and others have previously identified as particularly deregulated in ACC.^8, 25^

**Figure 2.**
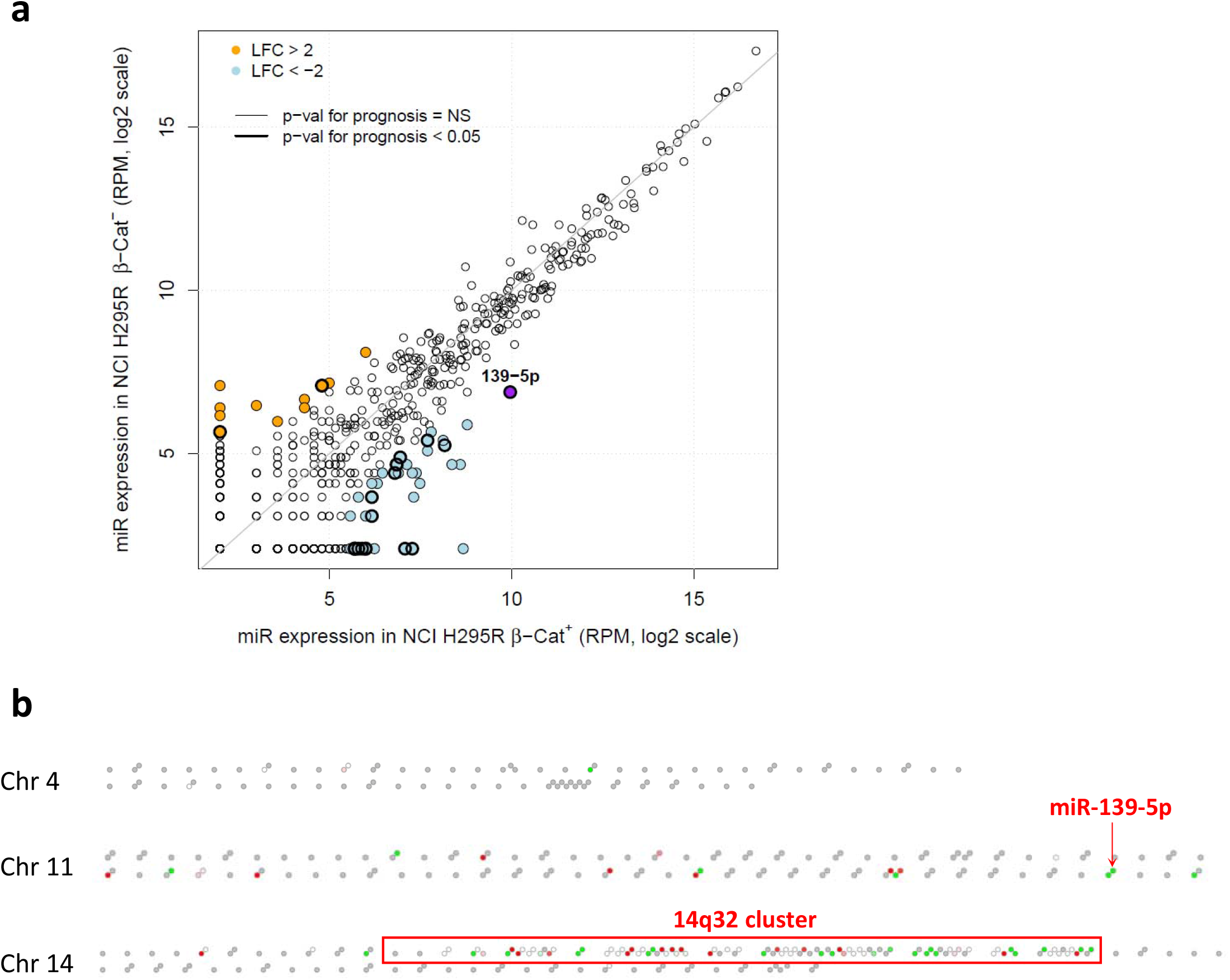
β-Catenin silencing impacts miRNA landscape in ACC. (a) Differential expression of miRNAs in H295R β-Cat^+^ versus H295R β-Cat^-^ cells. Light blue dots represent miRNAs that are downregulated (RPM > 10 in one condition and LFC < -2), orange dots represent miRNAs that are upregulated (RPM > 10 in one condition and LFC > 2) after β-Catenin silencing. Coloured dots with a thick border correspond to prognostic miRNAs (p<0.05, calculated for the COMETE-ENSAT cohort). NS: non-significant for prognosis. (b) Visualization of the genomic position of miRNAs whose expression is changed following β-Catenin silencing in H295R cells. To better visualize the global landscape of miRNAs, the selection criteria were the following : log2 RPM > 4 and LFC either > 1 or < -1. Green dots represent miRNAs that are downregulated (LFC < -1) while red dots represent miRNAs that are upregulated (LFC > 1) in β-Cat^-^ cells. In chromosome 4, only one miRNA was decreased after β-Catenin silencing. In chromosome 11, few miRNAs were changed, notably miR-139-5p. By contrast, in chromosome 14, numerous miRNAs clustered within in the 14q32 locus were affected.

Among the miRNAs that we found affected by β-Catenin silencing, we selected 8 of them for RT-qPCR validation, based on their p-value for prognosis (Supplementary Figure S3), including 4 miRNAs of the 14q32 locus (miR-377-5p, miR-377-3p, miR-1185-2-3p and miR-541-3p), miR-139-5p (chr 11), miR-320b (chr 1), miR-2110 (chr 10), and miR-513a-5p (chr X). In addition, we also measured the expression of miR-483-5p and miR-483-3p, two well-recognized miRNA markers of malignancy.^19^ As the expression of miR-1185-2-3p was below the detection limit (Ct > 35), this miRNA was not further analyzed. Reducing β-Catenin protein level over time led to a progressive and significant decrease of miR-139-5p expression (Figure 3a), with a 94 ± 0.1 %-decrease at D10. This decrease was not observed in control pTer1 cells (Supplementary Figure S4). In agreements with the RNA sequencing data, no changes were observed in the expression of miR-483-5p (Figure 3b) and miR-483-3p (Supplementary Figure S5). For all other selected miRNAs, we confirmed a significant decrease in their expression ranging from 35 % to 75 % at D10 of β-Catenin silencing (Supplementary Figure S6).

**Figure 3.**
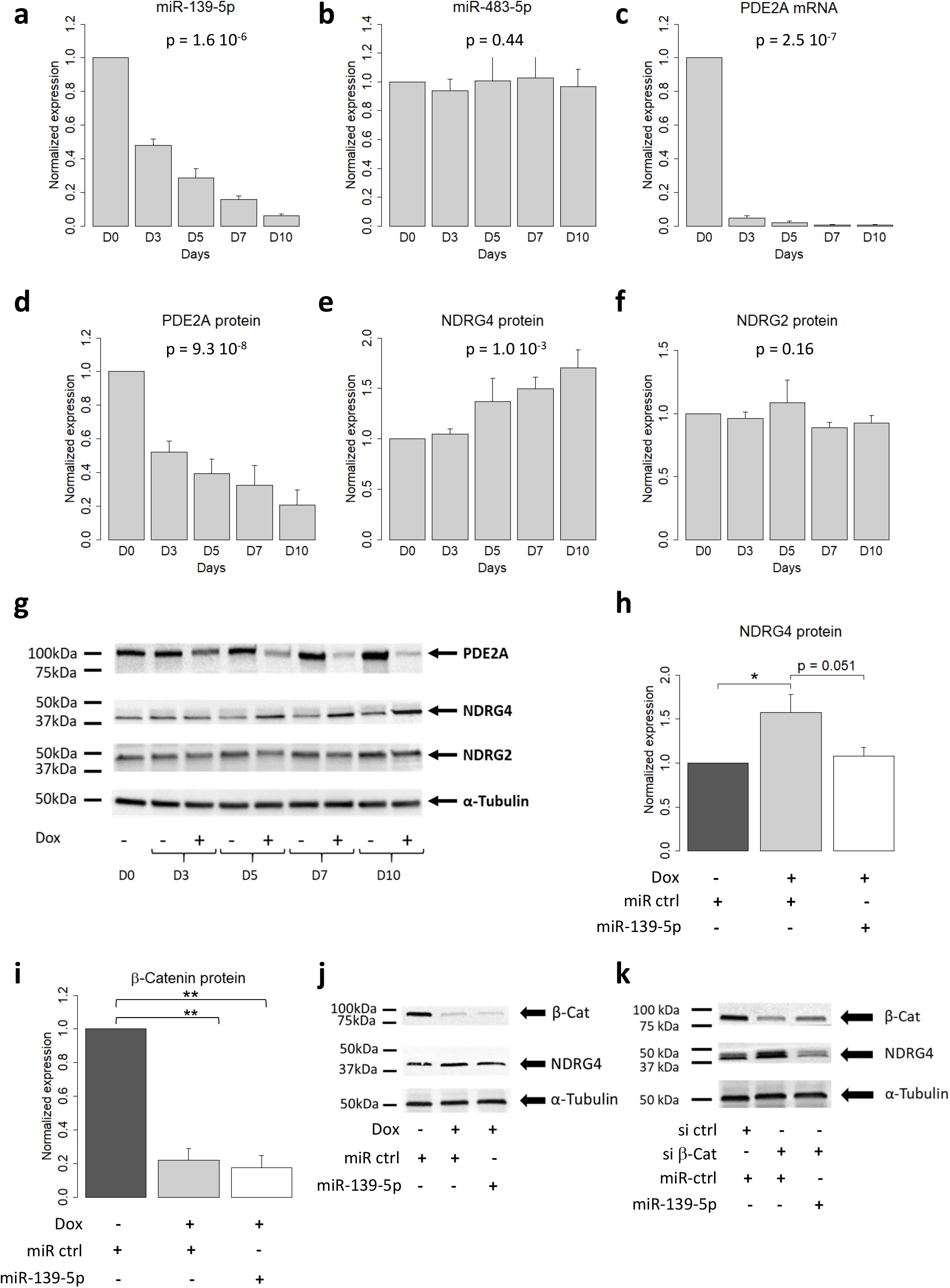
β-Catenin modulates the miR-139-5p/PDE2A/NDRG4 axis. (a), (c) and (d) β-Catenin repression progressively and markedly decreases the expression of miR-139-5p and its host gene PDE2A while it upregulates miR-139-5p target gene NDRG4 (e). The levels of miR-139-5p, miR-483-5p and PDE2A mRNA were assessed using RT-qPCR in H295R β-Cat^-^ cells. (b) and (f) Neither miR-483-5p nor its target gene NDRG2 were affected after β-Catenin silencing. (g) A representative western blot for PDE2A, NDRG4, and NDRG2 with α-tubulin as a loading control. Graphs for qPCR represent the mean ± S.E.M of 5 independent experiments performed in duplicate. Graphs for western blots represent the mean ± S.E.M of 4 independent experiments for PDE2A and 5 independent experiments for NDRG4 and NDRG2. (h) Rescuing miR-139-5p in H295R β-Cat^-^ cells prevents the increase in NDRG4 protein observed at D10 after b-Cat repression but does not change the expression of β-Catenin protein itself (i). Graphs for rescue experiments represent the mean ± S.E.M of 3 independent experiments. (j, k) A representative western blot of β-Catenin and NDRG4 in miR-139-5p rescue experiments using shβ7 cells treated or not with Doxycycline, or H295R cells transfected with control or β-Catenin siRNA, respectively. α-tubulin was used as a loading control.

### Wnt/β-Catenin signaling exerts a tight control over miR-139-5p, PDE2A and NDRG4 expression

The striking effect of β-Catenin silencing on miR-139-5p expression led us to focus on this miRNA. MiR-139-5p is an intronic miRNA, which is located in the second intron of Phosphodiesterase 2A (*PDE2A*). We thus investigated whether the decrease in miR-139-5p expression following β-Catenin silencing was correlated with a decrease in the expression of its host gene. As shown in Figure 3c, repression of β-Catenin led to a sharp and significant decrease of more than 90 ± 2 % in the expression of *PDE2A* mRNA, which persisted over time. This was accompanied by a progressive decline in PDE2A protein expression from 48 ± 7 % at D3 up to 79 ± 8 % at D10 (Figure 3d). PDE2A mRNA and protein expression levels were not affected in the control pTer1 cells (Supplementary Figure S7).

We have previously shown that NDRG4 is a direct target gene of miR-139-5p in H295R cells and that inhibition of miR-139-5p induces a significant increase in NDRG4 expression.^26^ Moreover, the levels of NDRG4 mRNA and protein were inversely correlated to those of miR-139-5p in ACC patients. We then investigated the link between β-Catenin-mediated regulation of miR-139-5p and its target gene *NDRG4*. Unexpectedly, the levels of *NDRG4* mRNA were significantly decreased in β-Cat^-^ cells (Supplementary Figure S8). By contrast, NDRG4 protein levels were progressively increased (Figure 3e and 3g) and were inversely correlated to those of miR-139-5p. NDRG4 mRNA and protein levels were unchanged in control pTer1 cells (Supplementary Figure S9). These results suggest that downregulation of miR-139-5p following β-Catenin repression is accompanied by a relief of miR-139-5p-mediated translational repression of NDRG4 mRNA. Interestingly, we found that the levels of NDRG2 were unchanged in β-Cat^-^ cells as observed for its regulatory miRNA miR-483-5p (Figure 3b, 3f and 3g). This result points at a specific effect of β-Catenin on miR-139-5p/NDRG4 axis.

To confirm the results obtained with the inducible β-Catenin shRNA, we used an siRNA-based strategy to silence β-Catenin expression transiently (Supplementary Figure S10). In H295R cells transfected with a β-Catenin siRNA, β-Catenin mRNA and protein expressions were significantly decreased by 57.2 ± 6% (p = 0.024, n = 3) and 60 ± 5 % (p = 0.001, n = 3), respectively, at D10 (Supplementary Figure S10a and 10e). *AXIN2* mRNA was decreased by 49.3 ± 8% (p = 0.025, n = 3) while protein expression was reduced by 61.3 ± 2% (p < 0.001, n = 3) at D10 (Supplementary Figure S10b and 10f). We also confirmed a decrease of *PDE2A* mRNA by 66.1 ± 1% (p = 0.008, n = 3) and PDE2A protein by 52.0 ± 5% (p = 0.002, n = 3, Supplementary Figure S10c and 10g). MiR-139-5p expression was decreased by 74 ± 2% (p < 0.001, n = 3) at D10 with a concomitant increase in NDRG4 protein level by 1.56 ± 0.08 fold (p = 0.006, n = 3) (Supplementary Figure S10j and 10h). Consistent with our previous findings, NDRG4 mRNA expression level was decreased (Supplementary Figure S10d) and the levels of miR-483-5p and miR-483-3p were unchanged at D10 upon siRNA-mediated β-Catenin repression (Supplementary Figure S10k and 10l). These data confirm that β-Catenin siRNA produces similar effects to those observed with β-Catenin shRNA.

To better assess the link between miR-139-5p, NDRG4 and β-Catenin, we conducted rescue experiments by transfecting exogenous miR-139-5p mimics in β-Cat^-^ cells. In β-Cat^-^ cells transfected with control miRNA mimics, NDRG4 protein level was 1.66 ± 0.27 higher than that of β-Cat^+^ cells (Figure 3h and 3j). Interestingly, adding exogenous miR-139-5p in β-Cat^-^ cells prevented the increase in NDRG4 protein level (1.10 ± 0.13-fold relative to control β-Cat^+^ cells, (Figure 3h and 3j). Similar results were obtained in H295R cells transfected with β-Catenin siRNA and control miRNA mimics or miR-139-5p mimics. Indeed, co-transfection of β-Catenin siRNA and miR-139-5p mimic prevented NDRG4 protein upregulation and even decreased the levels of NDRG4 detected in control β-Cat^+^ cells by 47.5 ± 0.02 % (Figure 3k and Supplementary Figure S11a). Interestingly, miR-139-5p mimics did not impact the levels of β-Catenin protein itself in both β-Cat^-^ and H295R cells transfected with β-Catenin siRNA (Figure 3i, 3j, 3k and Supplementary Figure S11b). Moreover, transfection of H295R cells with antimiR-139-5p showed no variations in either TCF-driven luciferase activity or β-Catenin protein level (Supplementary Figure S11c, 11d and 11e), thus ruling out a potential feedback loop between miR-139-5p and mutated β-Catenin in ACC cells. Altogether, these results suggest a β-Catenin-mediated control of miR-139-5p and NDRG4 expressions in H295R cells.

ACC cell models have been lacking for many years until the recent development of MUC-1 cell line, which has been generated from an ACC neck metastasis of a male patient.^31^ To explore more deeply the relationship between miR-139-5p and β-Catenin, we also used MUC1 cells, which express the wild type β-Catenin gene. We found that β-Catenin mRNA and protein expression levels in MUC1 cells represent respectively 40 ± 5% (p < 0.01, n = 3, Figure 4a) and 11 ± 2.6% of those of H295R cells (p < 0.001, n = 3, Figure 4b and 4i). Axin2 mRNA and protein levels were also lower in MUC1 cells, with an mRNA expression level of 3 ± 0.1% (p < 0.001, n = 3, Figure 4c) and protein level of 2 ± 0.3% of the H295R cell expression levels (p < 0.001, n = 3, Figure 4d and 4i). PDE2A mRNA and protein expression levels were barely detectable in MUC1 compared to H295R cells with an mRNA level representing 0.01 ± 0.04% (p < 0.001, n = 3, Figure 4e) and a protein level representing 3.6 ± 0.2% (p < 0.001, n = 3, Figure 4f and 4i) of H295R cell expression levels. MiR-139-5p expression was also sharply lower in MUC1 cells and represents 0.01 ± 0.1% of miR-139-5p levels in H295R cells (p < 0.001, n = 3, Figure 4g). Conversely, NDRG4 protein expression level was 5.7 ± 1.0-fold higher in MUC1 cells compared to H295R cells (p < 0.05, n = 3, Figure 4h and 4i).

**Figure 4.**
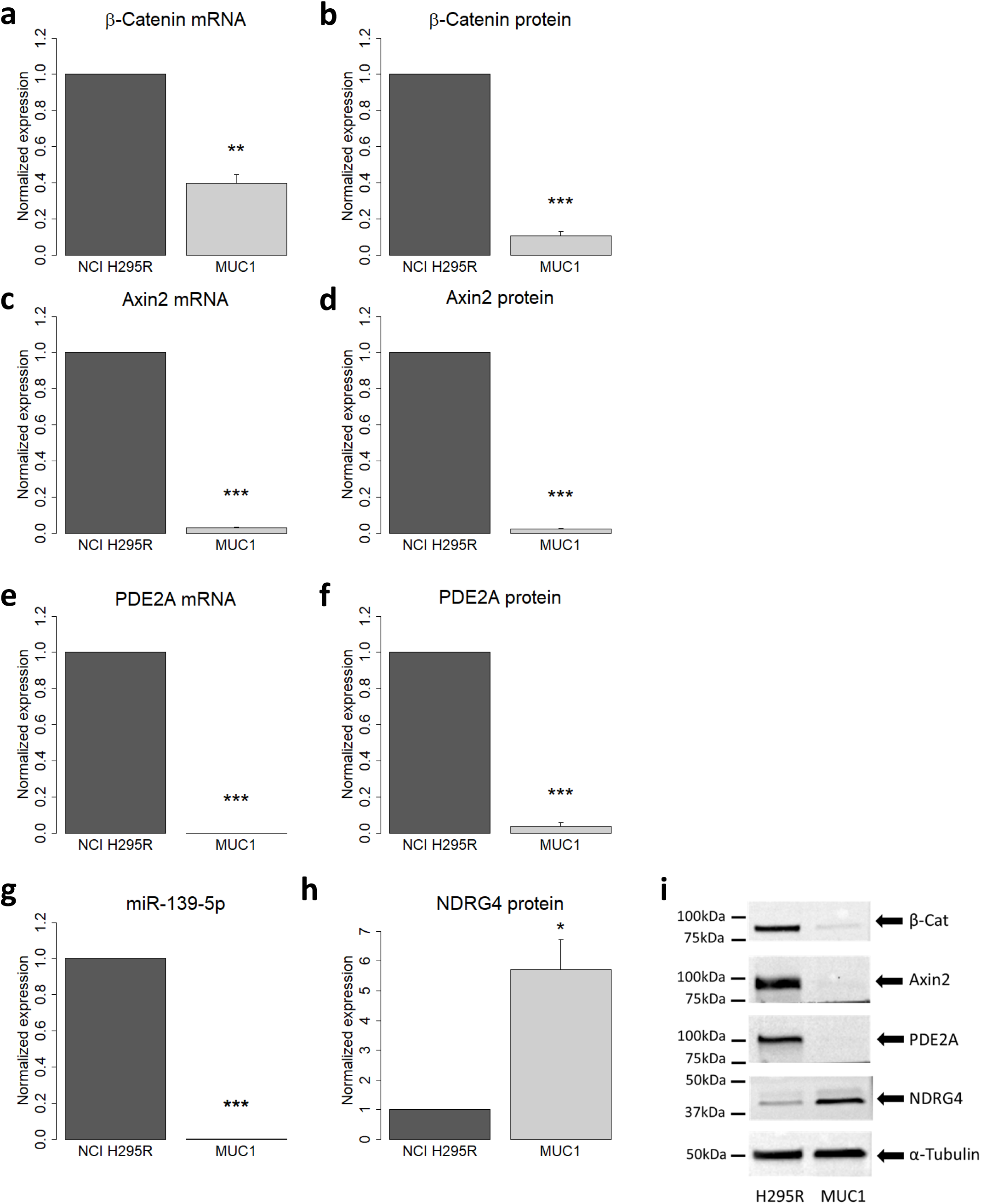
Comparison of selected mRNA and protein abundances in MUC1 and H295R cells. Shown are the expression levels of (a) β-Catenin mRNA, (b) β-Catenin protein, (c) Axin2 mRNA, (d) Axin2 protein, (e) PDE2A mRNA, (f) PDE2A protein, (g) miR-139-5p and (h) NDRG4 protein. Graphs show the mean ± S.E.M of 3 independent experiments. (i) A representative western blot analysis of the different proteins.

### β-Catenin gain of function and LEF1 upregulation are associated with increased miR-139-5p and PDE2A expression and reduced NDRG4 expression in ACC patients

To determine the clinical significance of our findings, we investigated the level of miR-139-5p and PDE2A in patients depending on the presence of genetic alterations in the Wnt/β-Catenin signaling pathway, as defined in Materials and Methods. In the COMETE-ENSAT cohort, patients with mutations in the Wnt/β-Catenin signaling pathway had significantly higher levels of miR-139-5p and PDE2A compared with patients without mutations (median expression of miR-139-5p: 12.8 vs 9.7, p = 0.015; median expression of PDE2A: 6.42 vs 4.57, p = 0.010, Figure 5a and 5b), and significantly lower levels of NDRG4 (median expression of 5.49 vs 6.65, p = 0.008, Figure 5c). Interestingly, the levels of PDE2A and miR-139-5p were strongly correlated in both COMETE-ENSAT (correlation coefficient: 0.94, p-value: 9.2 × 10^−21^) and TCGA (correlation coefficient: 0.96, p-value: 2 × 10^−42^) cohorts (Figure 5d and 5e), suggesting a co-transcriptional regulation of miR-139-5p and its host gene PDE2A in the tumor as observed in H295R cells in vitro.

**Figure 5.**
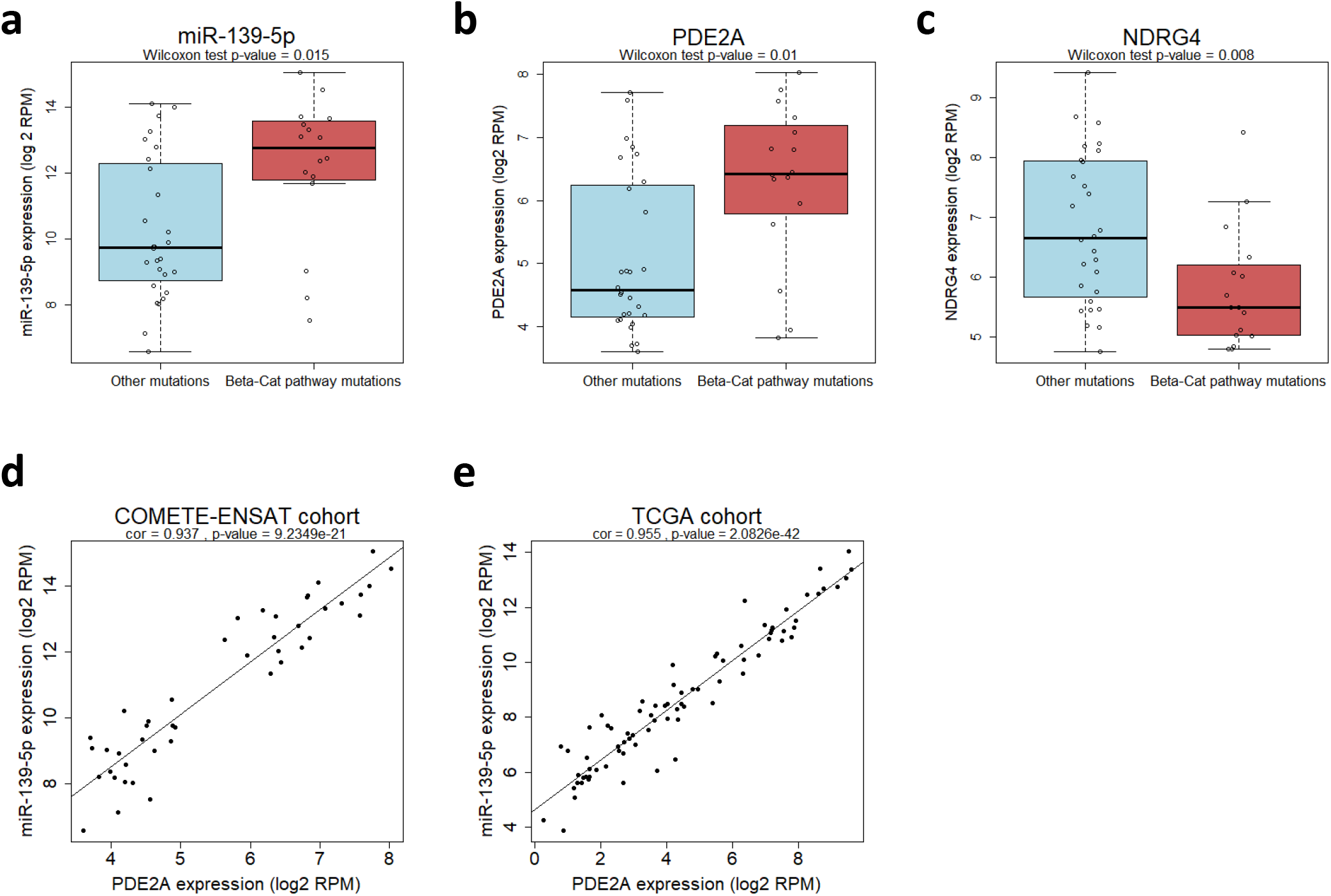
miR-139-5p and its host gene PDE2A are upregulated and strongly correlated in ACC patients with mutations in Wnt/β-Cat pathway. (a), (b) and (c) Expression of miR-139-5p, its host gene PDE2A and its target gene NDRG4 in the COMETE-ENSAT cohort of ACC patients, depending on the presence or the absence of mutations in Wnt/β-Catenin pathway. Patients with mutations in *CTNNB1* gene (β-Cat), *ZNRF3* gene or mutation in either gene in addition to *TP53* were listed as mutated for the Wnt/β-Catenin signaling pathway. Patients with mutations of *TP53, CDKN2A, MEN1, RB1, TERT, DAXX*, or *MED12* genes were listed as non-mutated for Wnt/β-Catenin signaling pathway (other mutations). (d) and (e) Tumor expression of miR-139-5p with respect to its host gene PDE2A in the COMETE-ENSAT cohort and the TCGA cohort. Each dot corresponds to a patient.

However, we discovered that a sub-group of 7/28 patients (25%) in the COMETE-ENSAT cohort had no identified mutation in the Wnt/β-Catenin signaling pathway but displayed a high level of miR-139-5p while, on the contrary, 3/16 patients (19%) carried a mutation in Wnt/β-Catenin signaling pathway and had a low level of miR-139-5p. We thus investigated whether the activation of Wnt/β-Catenin signaling pathway could be better predicted by the expression level of the TCF/LEF transcription factors than by the mutational status of β-Catenin. In vertebrates, TCF/LEF family includes four members: TCF1 (also called TCF7), LEF1 (or TCF1α), TCF3 and TCF4.^40^ We therefore analyzed the distribution of the expression of these 4 genes among patients. We found that LEF1 was the only transcription factor whose expression pattern was bi-modal (Supplementary Figure S12) with a local minimum around 5.2. Almost all β-Catenin-mutated patients have an expression level of *LEF1* above this threshold of 5.2, but interestingly, 12/28 (42.9 %) of the patients with no known mutation in the β-Catenin pathway had a high expression of *LEF1*, while only 2/16 (12.5 %) of the mutated patients had a low expression of *LEF1*.

To determine whether miR-139-5p expression correlates better with LEF1 expression than with Wnt/β-Catenin pathway mutational status, we examined the level of miR-139-5p in function of the level of *LEF1*. Patients with a *LEF1* expression level above 5.2 were classified in the “*LEF1* high” group while patients with a *LEF1* expression below 5.2 were classified in the “*LEF1* low” group. As shown in Figure 6a, the level of miR-139-5p in “*LEF1* high” patients was significantly higher than in the “*LEF1* low” patients (median 12.63 vs 8.74, p = 1 × 10^−7^) in the COMETE-ENSAT cohort, with a much more significant p-value than that obtained with the mutational status of Wnt/β-Catenin signaling pathway. Interestingly, all the patients with a tumor expression of miR-139-5p above 11 are now in the “LEF1 high” group. This observation was confirmed by analyzing the TCGA dataset (Figure 6b): the median expression of miR-139-5p in “*LEF1* high” patient group was significantly higher than that of the “*LEF1* low” patient group (10.25 vs 6.78, p = 9.3 × 10^−10^, LEF1 threshold 3.3). Moreover, in both cohorts, miR-139-5p expression shows a clear correlation with *LEF1* expression (Figure 6c and 6d). Besides, we observed a higher expression of *PDE2A* (p = 3.59 × 10^−7^) and lower expression of NDRG4 (p = 7.02 × 10^−5^) in the “LEF1 high” group in comparison to the “LEF1 low” group in the COMETE-ENSAT cohort (Figure 6e and 6f) and the TCGA cohort (Supplementary Figure S13a and 13b). As observed for miR-139-5p, *LEF1* performs better than Wnt/β-Catenin signaling pathway mutational status in discriminating patients according to their *PDE2A* and *NDRG4* expression levels, in the COMETE-ENSAT and the TCGA cohorts.

**Figure 6.**
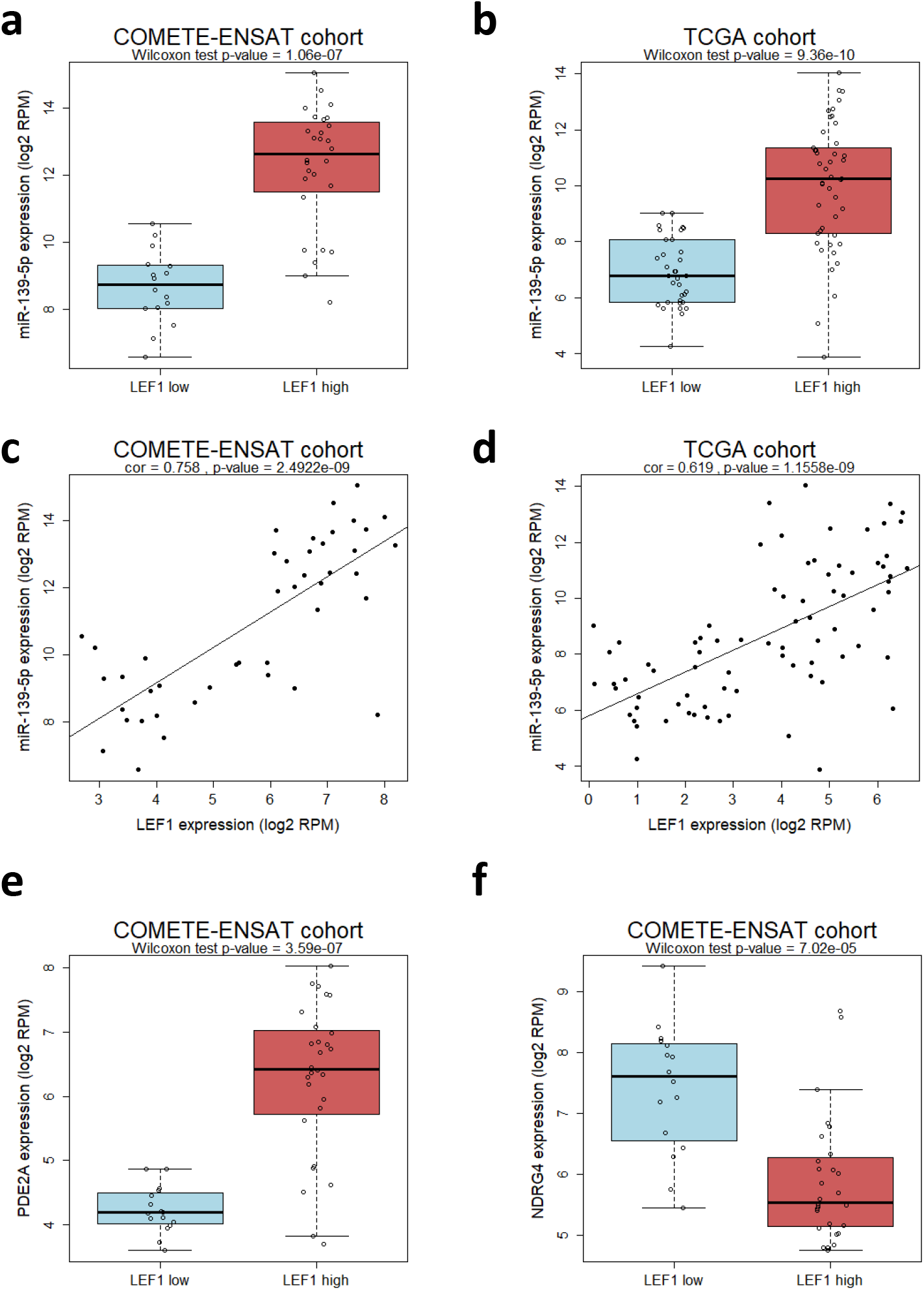
miR-139-5p and PDE2A expressions are tightly correlated with LEF1 expression levels in ACC patients. (a) and (b) Expression of miR-139-5p as a function of LEF1 expression level in the COMETE-ENSAT cohort and the TCGA cohort. The selected thresholds for LEF1 in the COMETE-ENSAT and the TCGA cohorts were 5.2 and 3.3, respectively, based on the Expectation-Maximization algorithm. (c) and (d) Expression of miR-139-5p as a function of LEF1 expression in both cohorts. (e) Expression of miR-139-5p host gene PDE2A and of its target gene NDRG4 (f) in the COMETE-ENSAT cohort, in function of LEF1 expression levels.

We have previously shown that miR-139-5p expression levels have a significant prognostic value for ACC.^26^ We thus evaluated the prognostic significance of miR-139-5p regulators in ACC patients, namely the mutational status of Wnt/β-Catenin signaling pathway and LEF1. Comparison of survival curves of patients in function of their mutational status with those in function of their LEF1 expression levels revealed that LEF1 better predicts overall survival than Wnt/β-Catenin pathway mutation status (Figure 7a and 7b, p = 0.018 and p = 0.0021 for the mutational status and LEF1 expression, respectively). The powerful predictive value of LEF1 expression was also confirmed in the TCGA cohort (p = 5.7 × 10^−5^, Figure 7c). Remarkably, prediction of survival using LEF1 closely mimics the prediction using the ACC C1A and C1B mRNA clusters determined from the entire transcriptome^8^ of the COMETE-ENSAT cohort (Figure 7d). Indeed, 2 death events were observed in both the “LEF low” group and the C1B group compared to 17 death events observed in both the “LEF high” group and the C1A group (p = 0.0021 for LEF1, and p = 0.0001 for C1A/C1B clusters).

**Figure 7.**
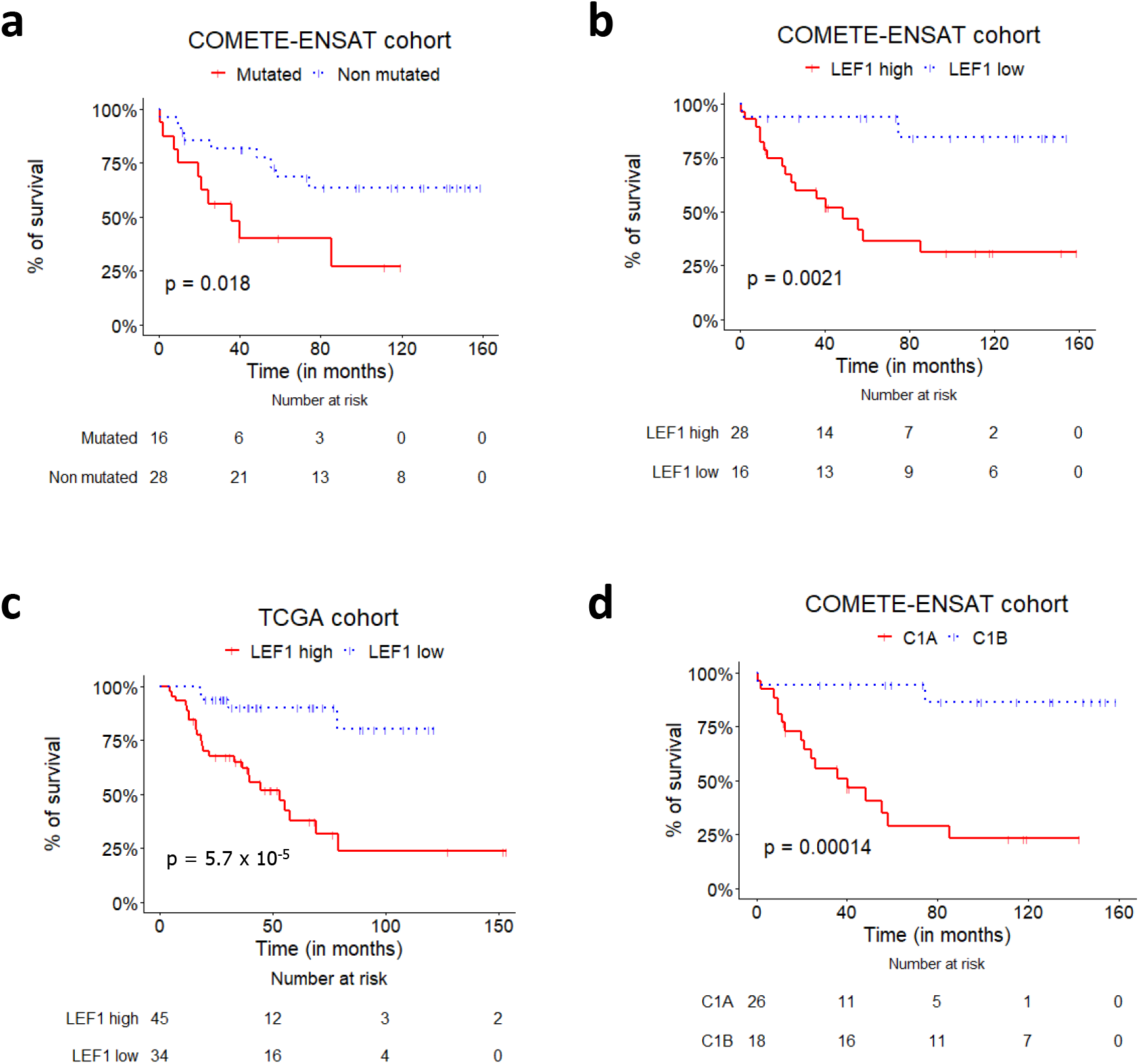
LEF1 is a better predictor of ACC patient outcome than Wnt/β-Cat pathway mutational status. (a) and (b) Overall survival in the COMETE-ENSAT cohort (n = 44) in function of the mutational status on Wnt/β-Catenin signaling pathway and LEF1 expression, respectively. (c) Overall survival in the TCGA cohort (n = 79) in function of LEF1 expression levels. (d) Overall survival in the COMETE-ENSAT cohort in function of the mRNA clusters C1A and C1B.

## Discussion

ACC is a rare malignancy with dismal prognosis and clinical heterogeneity. Understanding ACC biology is essential to develop new efficient therapies. Gain-of-function mutations and deletions in genes involved in the Wnt/β-Catenin signaling pathway have been reported in approximately 40% of ACC. Mutations in *CTNNB1* (β-Catenin) gene were found in 15-20% of the tumors,^8, 9^ with activating mutations in exon 3 impairing Serine/Threonine residue phosphorylations. Constitutive activation of Wnt/β-Catenin pathway in ACC is associated with poor prognosis.^13, 14, 41^ Besides, deregulations of tumor and circulating miRNAs were found in ACC patients and many of them showed tight correlation with prognosis. Nevertheless, the molecular mechanisms underlying aberrant expression of miRNAs in ACC are still unknown. As Wnt/β-Catenin signaling pathway is a key driver of both adrenal development and tumorigenesis, we hypothesized that this pathway may promote ACC aggressiveness through modulations of miRNA expression.

We report here for the first time that constitutively active Wnt/β-Catenin signaling contribute to miRNA expression deregulations in ACC cells. Indeed, RNA sequencing of the whole miRnome after inducible shRNA-mediated silencing of β-Catenin revealed that 54 miRNAs were affected (LFC>2 or <-2), mainly by a decrease in their expression (42 out of 54 miRNAs). A threshold of 2 for determining a significant change in gene expression (LFC) was chosen to select miRNAs that β-Catenin silencing impacted the most. However, when we used a less stringent threshold of 1, many other miRNAs were impacted (117 miRNAs including 85 with a decreased expression), suggesting a strong control of miRNA expression by β-Catenin. We subsequently confirmed by quantitative PCR that β-Catenin repression significantly decreases the expression of miR-377-5p, miR-377-3p, miR-541-3p (chr 14), miR-139-5p (chr 11), miR-2110 (chr 10) and miR-513a-5p (chr X), all of which have been found to be overexpressed in aggressive ACC and associated with prognosis.^8, 19^

Most studies of the relationship between miRNAs and Wnt/β-Catenin signaling pathway have focused on miRNAs as post-transcriptional regulators of β-Catenin expression. Notably, restoration of specific tumor suppressor miRNAs led to a decrease in β-Catenin mRNA expression through a direct binding of miRNAs to the 3’-untranslated region of β-Catenin transcript.^27-30^ To the best of our knowledge, about ten studies showed a direct effect of β-Catenin on miRNA expression in cancer.^42-50^ Ji *et al*. found that the Wnt/β-Catenin signaling pathway activates transcription of the miR-181s family (miR-181a, b, c and d) in hepatocellular carcinoma.^44^ Tang *et al*. showed that β-Catenin/TCF complex directly binds to the promoter of miR-183-96-182 cluster and activates its transcription in gastric cancer, leading to an increase in cell proliferation.^47^ *Guo et al*. reported a direct effect of β-Catenin on miR-150 transcription in colorectal cancer cells with subsequent increases in invasion, migration and epithelial-mesenchymal transition.^50^

Among the miRNAs whose expression varies following β-Catenin silencing, we identified a subset of both upregulated and downregulated miRNAs located in the 14q32 imprinted locus, which encodes for 54 miRNAs and is the largest miRNA cluster known in the human genome. Depending on the cancer type, miRNAs at this locus act as oncomiRs or tumor suppressor miRNAs, as reviewed by Benetatos *et al*.^51^ We first described the overexpression of 13 miRNAs at the 14q32 locus in aggressive ACC.^25^ Consistent with our data, an overall under-expression of this locus was found in the good prognosis subgroup of ACC.^8^ More recently, it was reported that deregulations of the 14q32 miRNA cluster in endocrine tumors could have a significant impact on prognosis.^52^ Interestingly, miR-410, miR-433 and miR-377-3p within the 14q32 locus have been identified as regulatory miRNAs of Wnt/β-Catenin signaling.^53, 54^ Conversely, we observed a significant decrease in miR-377-3p and miR-410 expression and slight up-or down-regulation of the majority of 14q32 miRNAs upon β-Catenin silencing in ACC cells. Altogether, these data suggest a feedback circuit between miRNAs and Wnt/β-Catenin pathway.

The pronounced effect of β-Catenin silencing on miR-139-5p expression together with the fact that we and others have previously identified miR-139-5p as a marker of aggressiveness and poor prognosis in ACC led us to focus on this miRNA. We demonstrate here *in vitro* that miR-139-5p expression is strongly decreased following β-Catenin silencing, in correlation with a complete repression of its host gene *PDE2A*. PDE2A is a dual-substrate phosphodiesterase that degrades both cGMP and cAMP. It is highly conserved among species and expressed in numerous organs in human, particularly in brain and endocrine organs including the adrenal, the pituitary, the pancreas and the mammary gland.^55^ The role of PDE2A in development is crucial since PDE2A-KO mice die between the second and the third week of gestation with major heart defects.^56^ PDE2A expression was found to be prominent in the zona glomerulosa (ZG) as compared to the zona fasciculata (ZF) of the adrenal gland. The maintenance of these two physiologically distinct zones would rely on the reciprocal inhibitory effects of the cAMP/Protein Kinase A and the Wnt/β-Catenin pathways in the ZF and the ZG, respectively.^57-60^ The importance of PDE2A in the maintenance of adrenal zonation has been reported recently. Indeed, Pignatti *et al*. identified PDE2A as a direct target of β-Catenin.^61^ They demonstrated that β-Catenin gain-of-function in the ZG induces an upregulation of PDE2A and impairs ZG-to-ZF cell transdifferentiation through inhibition of cAMP/PKA signaling in the ZF. Whether miR-139-5p also contributes to the identity of ZG remains to be determined. In adrenocortical tumors, Durand *et al*. first identified upregulation of *PDE2A* in adenomas and ACC harboring *CTNNB1* mutation.^62^ The parallel decrease in PDE2A and miR-139-5p levels that we observed in H295R cells *in vitro* suggests that the β-Catenin-dependent expression of miR-139-5p is coupled to that of its host gene *PDE2A*. Conversely, MUC1 cells, which originate from a neck ACC metastasis with no activating mutation in β-Catenin gene, display reduced levels of β-Catenin and barely detectable levels of PDE2A and miR-139-5p compared to H295R cells. MiR-139-5p has been shown to regulate Wnt/β-Catenin signaling pathway in cervical cancer^63^ and bladder cancer^64^ by targeting TCF4 and WNT1, respectively. In ACC, we did not observe a regulatory feedback loop between miR-139-5p and Wnt/β-Catenin signaling as shown by TOP-Flash reporter assays.

We found that miR-139-5p expression was strongly correlated with *PDE2A* expression in the two European and American cohorts of ACC patients (COMETE-ENSAT and TCGA). Interestingly, a previous study showed that miR-139-5p was correlated with PDE2A expression in 30/31 cancers of the TCGA consortium.^65^ However, no correlation between miR-139-5p/-3p and *PDE2A* expression was observed *in vitro* in acute myeloid leukemia (AML), in gastric cancer (GC) and colorectal cancer (CRC), suggesting that depending on the cellular context, independent transcription start sites or post-transcriptional mechanisms could be involved.^66-68^ It is worth mentioning that, by contrast to ACC, miR-139-5p is underexpressed in AML, GC and CRC and acts as a tumour-suppressor miRNA in many cancers.^69, 70^

To strengthen our observations *in vitro*, we analyzed the relationship between Wnt/β-Catenin pathway mutational status and miR-139-5p levels in ACC patients. Importantly, we found that patients with mutations have significantly higher levels of *PDE2A* and miR-139-5p and a significantly lower level of *NDRG4* when compared to patients with no known mutation in the Wnt/β-Catenin pathway. However, we also found that 7 patients out of 28 had high levels of miR-139-5p in the absence of identified mutations Wnt/β-Catenin pathway. A potential mechanism that may explain these findings is a nuclear expression of active β-Catenin in some patients with wild type *CTNNB1*, as it has been reported previously.^41^ Furthermore, the nuclear localization of β-Catenin was shown recently to be promoted by LEF-1 in cancer cells.^71^ These findings led us to analyze miR-139-5p expression in function of the expression of LEF1, a well-established downstream target gene of β-Catenin in ACC.^8, 9, 41, 72^ We postulated that the level of LEF1 transcription factor could reflect the global activity of Wnt/β-Catenin pathway in ACC whether it holds a known mutation or not. Remarkably, LEF1 was the only transcription factor of the LEF/TCF family members tested, whose expression tightly correlates with that of miR-139-5p. LEF1 better separates patients with high and low expression of miR-139-5p, PDE2A and NDRG4 than does the β-Catenin mutational status. We also identified LEF1 as a powerful survival predictor in ACC, with more statistical power than the mutational status of Wnt/β-Catenin pathway. In addition, survival prediction with LEF1 is comparable to that obtained with the whole transcriptome defining the C1A and C1B ACC subgroups. Establishing prognosis with a single gene expression might be more practical in a translational perspective. These data confirm that transcript expression better predicts patients’ outcome than the mutational status, as recently reported in a genome-wide analysis of prognostic features in human cancers.^73^

In summary, our study provides the first analysis of the impact of aberrant activation of Wnt/β-Catenin signaling pathway on the whole miRnome in cancer cells. We demonstrate that β-Catenin is involved in the upregulation of several miRNAs which are associated with poor outcome in ACC, including those of the 14q32 cluster and most of all, miR-139-5p, that we have previously found upregulated in aggressive tumors. Expression of miR-139-5p is strongly correlated with the expression of its host gene PDE2A in ACC cells *in vitro* as well as in patients. Moreover, ACC with mutated Wnt/β-Catenin pathway display higher levels of miR-139-5p as compared to non-mutated ACC. Finally, we provide evidence that high LEF1 expression in ACC correlates even better with high miR-139 expression than the mutational status of Wnt//β-Catenin pathway and is associated with significantly worse survival for the patients. These findings may trigger further mechanistic studies on the role of Wnt/β-Catenin signaling in promoting ACC aggressiveness through miRNA-mediated circuits. Considering that a single aberrant miRNA can regulate entire gene networks, thereby substantially contributing to ACC development and progression, strategies targeting β-Catenin activity are worth exploring for therapeutic opportunities in ACC.

## Contributors

NC conceived and supervised the study. JC conducted the experiments and carried out RNA-seq data analysis and interpretation. LG helped with RNA-Seq analysis, clinical data acquisition, statistical analyses, and interpretation. JD provided technical assistance. SRS and JD performed experiments. NC and JC drafted the manuscript. BR, CH and OC reviewed the manuscript. All the authors read and approved the final manuscript.

## Acknowledgments

The authors would like to thank Christophe Battail from the U1292 INSERM lab (Grenoble, France) and Anne Boland-Auge from the Centre National de Recherche en Génomique Humaine (Paris, France) for helpful advices on RNA-sequencing. Funding for this study was provided by the Institut National de la Santé et de la Recherche Médicale, the Ligue Contre le Cancer (Comité Isère), the French Endocrine Society (SFE), and the Fonds AGIR pour les Maladies Chroniques (Grenoble, France).

## Data sharing statement

RNA sequencing data collected for the study, including raw data and data analysis will be made available to others upon request. All data will be available upon publication of the manuscript, by contacting the corresponding authors.

