## Supplemental Data for "Aberrant activation of Wnt/β-Catenin signaling pathway drives the expression of poor prognosis-associated microRNAs in adrenocortical cancer with a major impact on miR-139-5p and its host gene PDE2A"

### Supplementary Figures

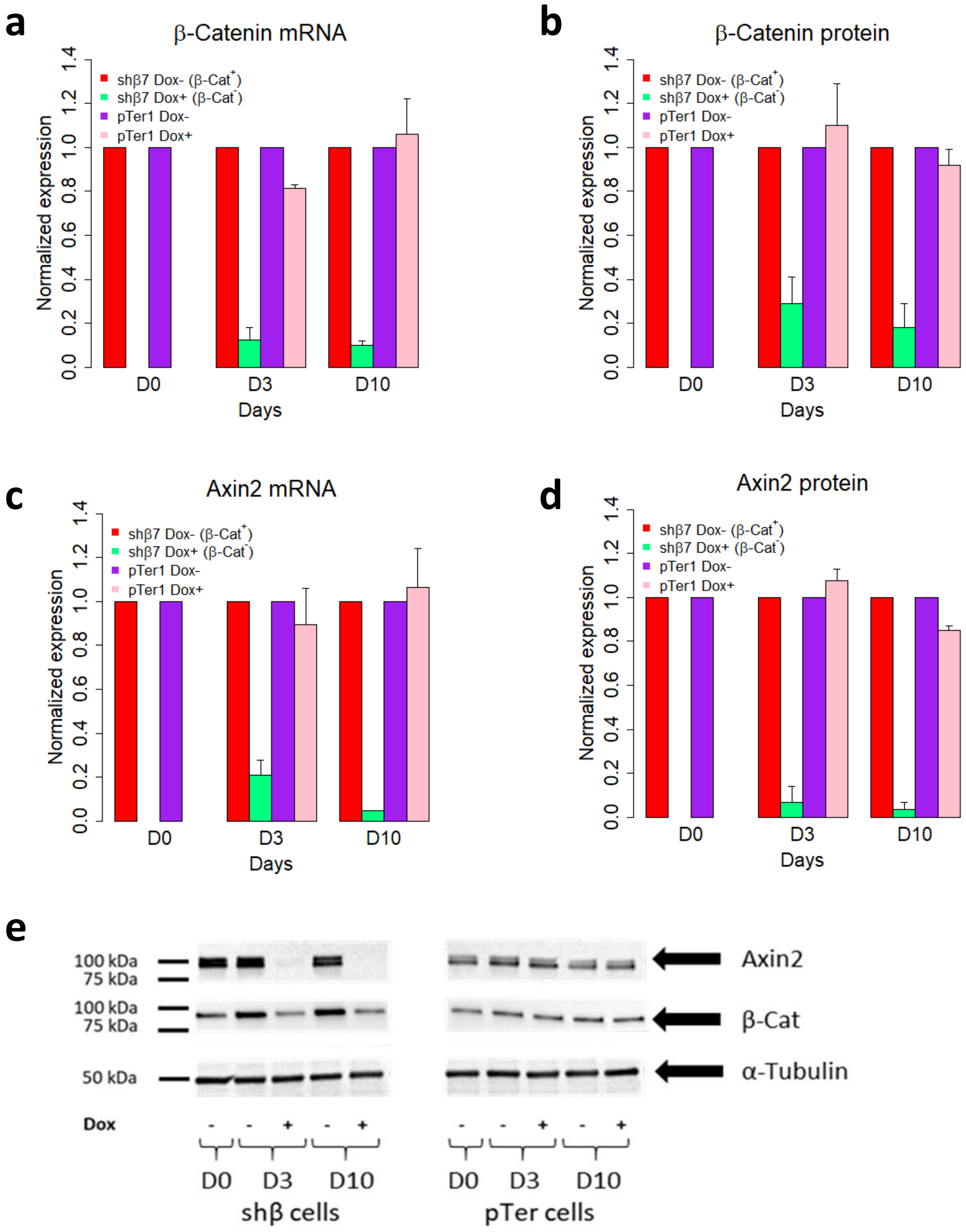


**Supplementary Figure S1. Doxycycline does not silence β-Catenin and Axin2 expression in H295R pTer1 control cells.** H295R shβ7 and pTer1 cells were cultured without (β-Cat+ cells) or with Doxycycline (β-Cat-) cells during 10 days. Cells were lysed at the day (D) indicated (D0, D3 and D10). (a) and (b) represent the levels of expression of β-Catenin mRNA and protein assessed by RT-qPCR and western blot, respectively. (c) and (d) represent the levels of expression of Axin2 mRNA and protein assessed by RT-qPCR and western blot, respectively. (e) A representative western blot for β-Catenin and Axin2 protein levels, with α-tubulin as a loading control. Graphs for qPCR and western blot analyses represent the mean ± SD of two independent experiments.

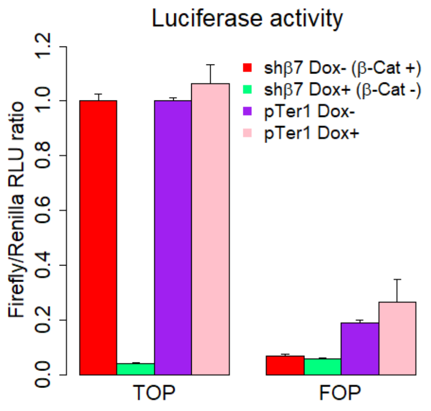


**Supplementary Figure S2. Doxycycline does not affect the TOP-FOP Flash luciferase activity in pTer1 control cells.** The graph represents the Firefly/Renilla activity ratio after 48h of culture with or without Doxycycline. Shown are the mean ± SEM of three experiments.

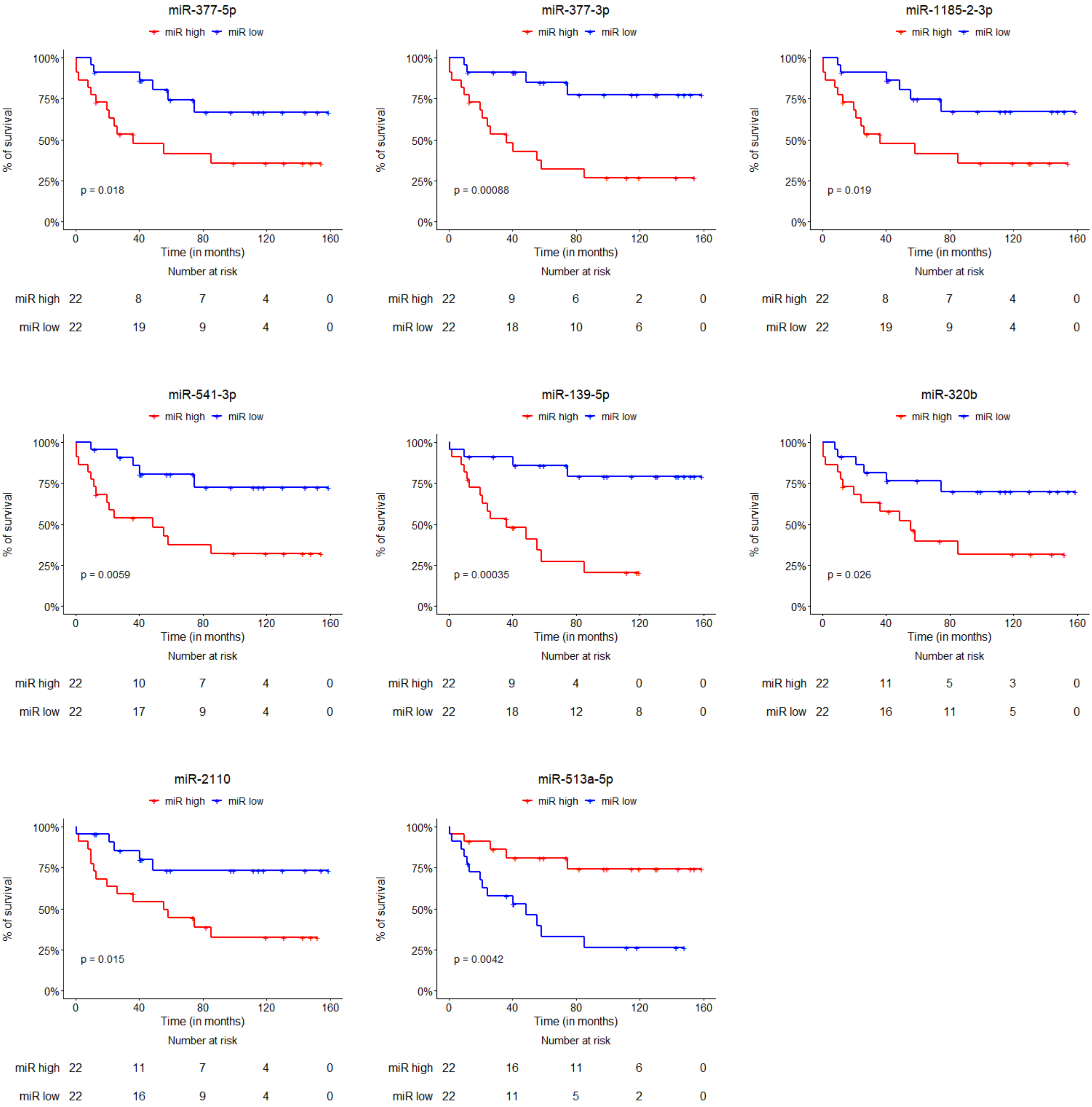


**Supplementary Figure S3. Overall survival of the patients of the COMETE-ENSAT cohort depending on miRNA expression in the tumor.** For each miRNA, the population was divided in two equal groups of 22 patients, the low and high groups corresponding to patients with a lower and higher expression as compared to the median value of the miRNA level. The p-value was calculated with the Log Rank test.

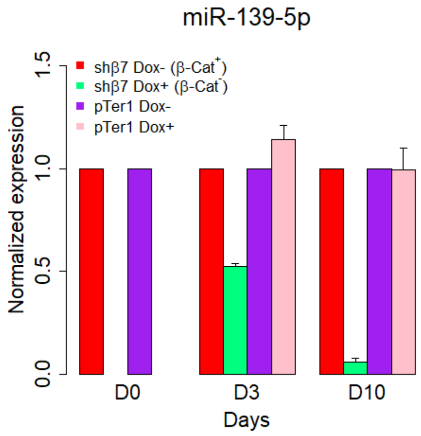


**Supplementary Figure S4. Doxycycline decreases miR-139-5p expression in H295R shβ7 cells but not in pTer1 control cells.** H295R shβ7 and pTer1 cells were cultured without (β-Cat$^{+}$ cells) or with Doxycycline (β-Cat$^{-}$) cells during 10 days. The graph shows the mean ± SD of two independent experiments.

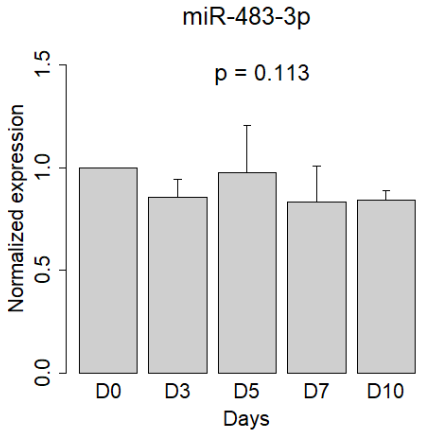


**Supplementary Figure S5. Silencing β-Catenin in H295R shβ7 cells does not affect miR-483-3p expression**. Cells were cultured without (β-Cat$^{+}$) or with Doxycycline (β-Cat$^{-}$) during 10 days. Cells were lysed at the day (D) indicated (D0, D3, D5, D7 and D10). Shown are RT-qPCR measurements of miR-483-3p levels in 3 independent experiments (mean ± SEM).

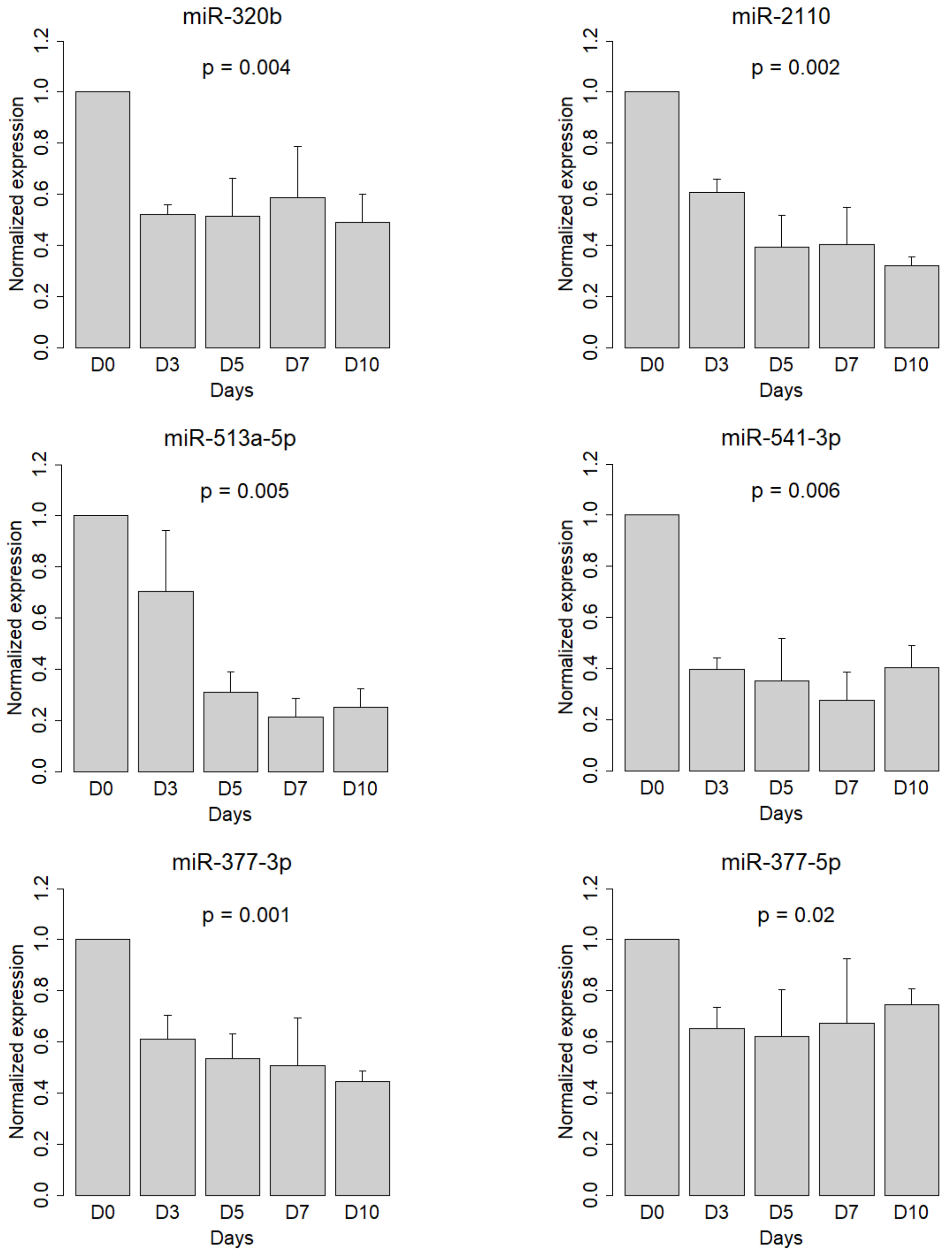


**Supplementary Figure S6. Validation of the decreased expression of selected miRNAs after silencing of β-Catenin in H295R cells.** Expression of miR-320b, miR-2110, miR-513a-5p, miR-541-3p, miR-377-3p and miR-377-5p were measured by RT-qPCR at the day (D) indicated (D0, D3, D5, D7 and D10) for β-Cat silencing duration. The graphs show the mean ± SEM of 3 independent experiments.

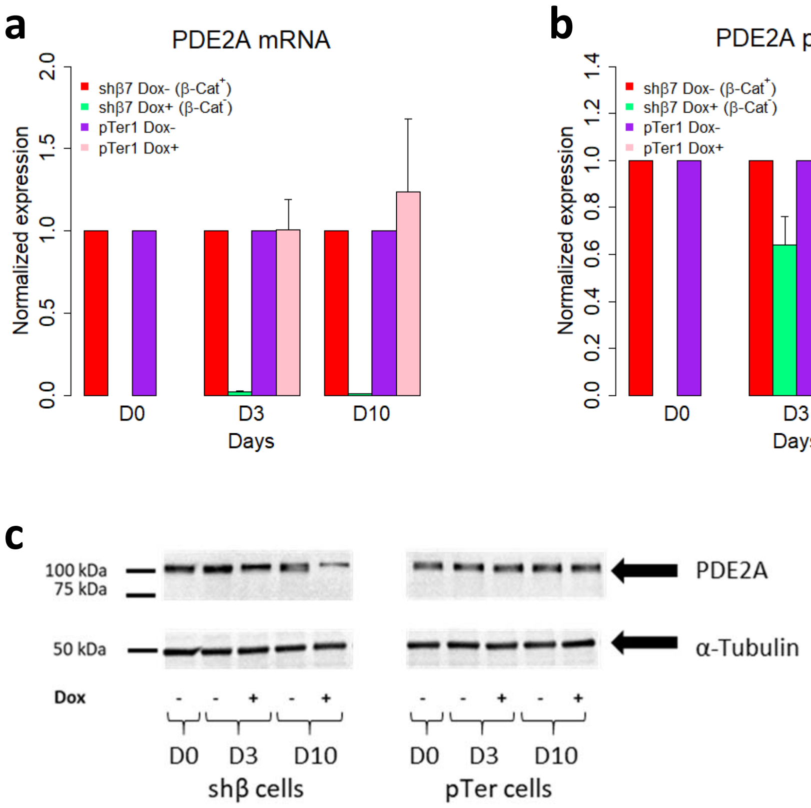


**Supplementary Figure S7. Doxycycline does not change PDE2A expression in H295R pTer1 control cells.** H295R shβ7 and pTer1 cells were cultured without (β-Cat+ cells) or with Doxycycline (β-Cat$^{-}$) cells during 3 or 10 days. Cells were lysed at the day (D) indicated (D0, D3 and D10). (a) and (b) represent the levels of expression of PDE2A mRNA assessed by RT-qPCR and PDE2A protein expression assessed by western blot. (c) A representative western blot for PDE2A with α-tubulin used as a loading control. Graphs for qPCR and western blot analyses represent the mean ± SD of 2 independent experiments.

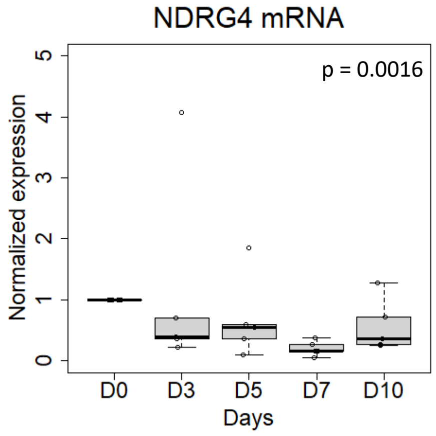


**Supplementary Figure S8. β-Catenin silencing in H295R shβ7 cells decreases NDRG4 mRNA expression.** Cells were cultured without (β-Cat$^{+}$) or with Doxycycline (β-Cat$^{-}$) during 10 days. Cells were lysed at the day (D) indicated (D0, D3, D5, D7 and D10). Shown are the median values of NDRG4 mRNA levels in 5 independent experiments.

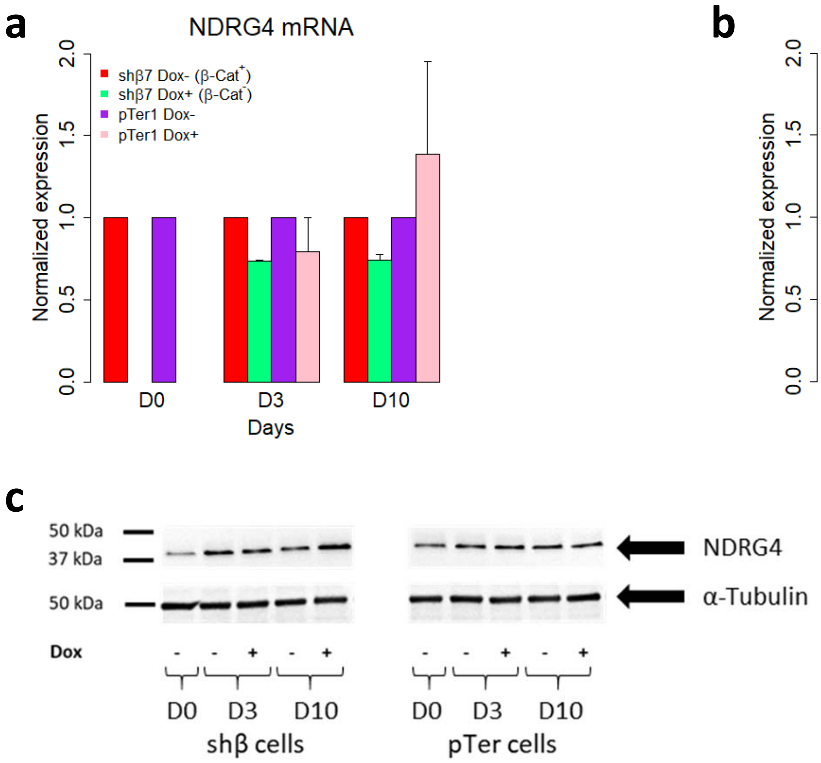


**Supplementary Figure S9. Doxycycline does not change NDRG4 mRNA and protein expression in H295R pTer1 control cells.** (a) and (b) Levels of expression of NDRG4 mRNA and protein in shβ7 and pTer1 cells treated or not with Doxycycline (mean ± SD of 2 independent experiments). (c) A representative western blot for NDRG4 in shβ7 and pTer1 cells.

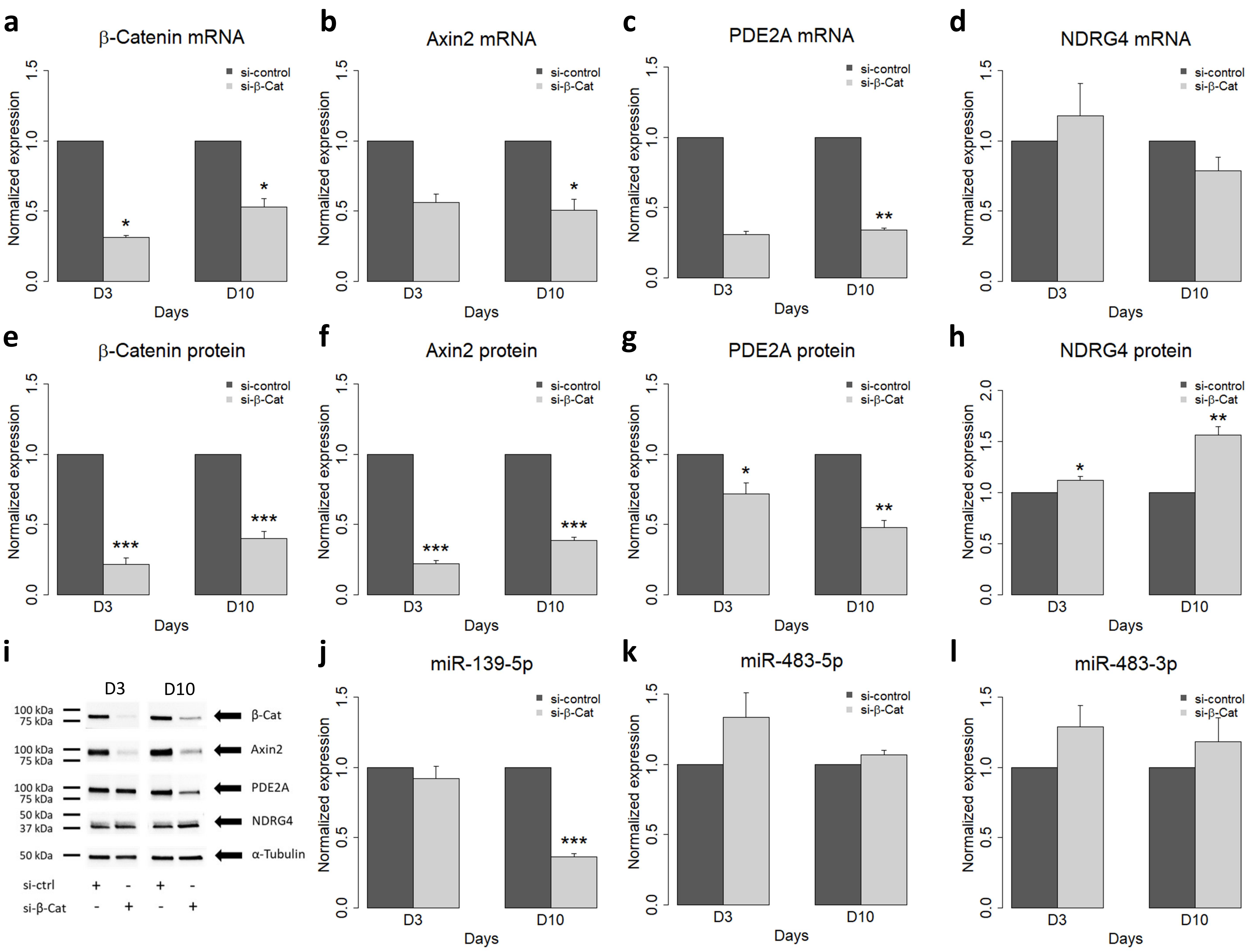
a
β-Catenin mRNA
b
Axin2 mRNA
c
PDE2A mRNA
d
NDRG4 mRNA
e
β-Catenin protein
f
Axin2 protein
g
PDE2A protein
h
NDRG4 protein
i
j
miR-139-5p
k
miR-483-5p
l
miR-483-3p
Normalized expression
Days
D3
D10
si-control
si-β-Cat
100 kDa
75 kDa
50 kDa
37 kDa
β-Cat
Axin2
PDE2A
NDRG4
α-Tubulin
si-ctrl

**Supplementary Figure S10. Silencing of β-Catenin in H295R cells using a specific siRNA represses Axin 2, PDE2A as well as miR-139-5p expression and upregulates NDRG4 protein levels.** Cells were transfected either with control siRNA or β-Cat siRNA at D0. The graphs represent the mean ± SEM in 3 independent experiments at D3 and D10 for (a) β-Cat mRNA, (b) Axin2 mRNA, (c) PDE2A mRNA, (d) NDRG4 mRNA, (e) β-Cat protein, (f) Axin2 protein, (g) PDE2A protein, (h) NDRG4 protein, (j) miR-139-5p, (k) miR-483-5p and (l) miR-483-3p. Statistical analyses are described in details in Materials and Methods. (i) shows a representative western blot of the 3 independent experiments.

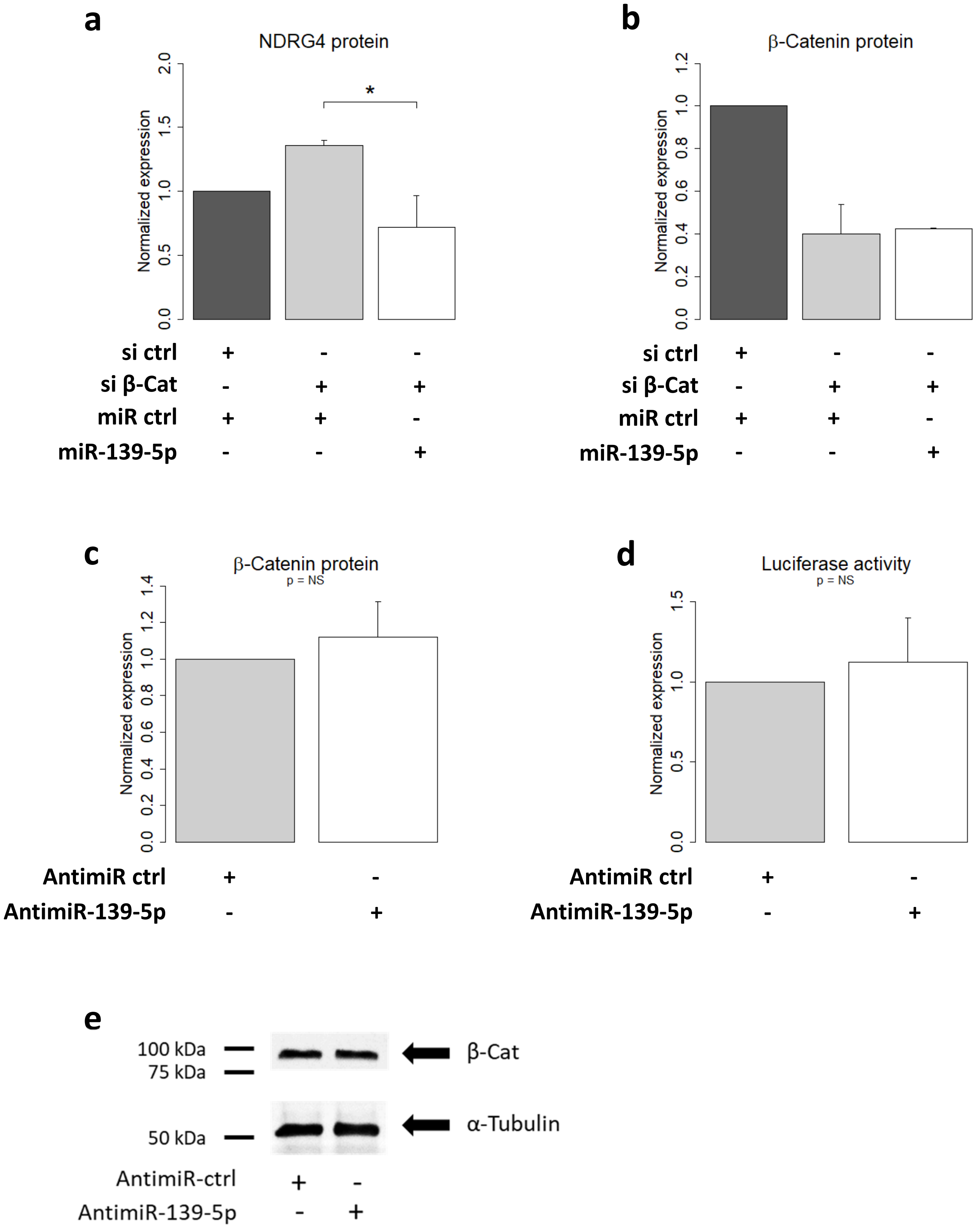


**Supplementary Figure S11. miR-139-5p modulates NDRG4 expression but does not affect Wnt/β-Catenin signalling.** (a) and (b) Rescuing miR-139-5p in H295R cells silenced for β-Cat using siRNA prevents the upregulation of NDRG4 protein but does not affect β-Cat levels. Graphs for rescue experiments represent the mean ± S.E.M of 2 independent experiments. (c) and (d) Inhibition of miR-139-5p expression with antimiRs in H295R cells does not change either the levels of β-Cat protein or β-Cat/TCF-driven luciferase activity. Shown are the mean ± SEM of three independent experiments. (e) A representative western blot for β-Cat with α-tubulin used as a loading control.

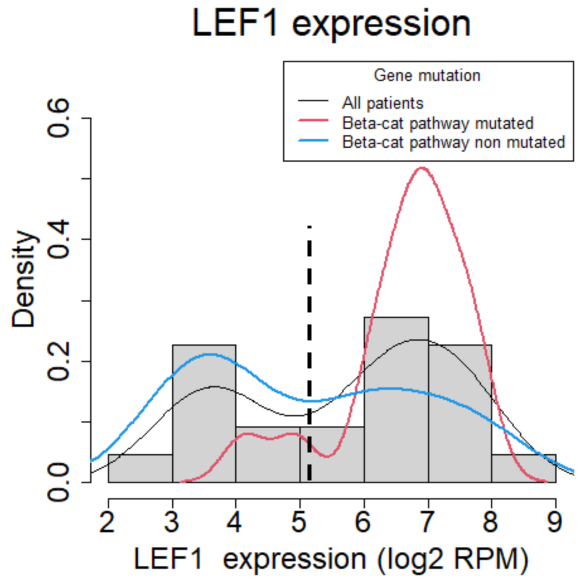


**Supplementary Figure S12. Distribution of LEF1 mRNA expression in the tumor of ACC patients from the COMET-ENSAT cohort.** The black line shows the bi-modal distribution of LEF1 expression in all the tumors, the blue line shows its distribution in tumors without known mutations in the Wnt/β-Cat pathway and the red line shows the distribution in tumors with known mutations in Wnt/β-Cat pathway. The vertical dotted line shows the local minimum at the threshold of 5.2 obtained from fitting Gaussian mixture with the Expectation-Maximization (EM) algorithm.

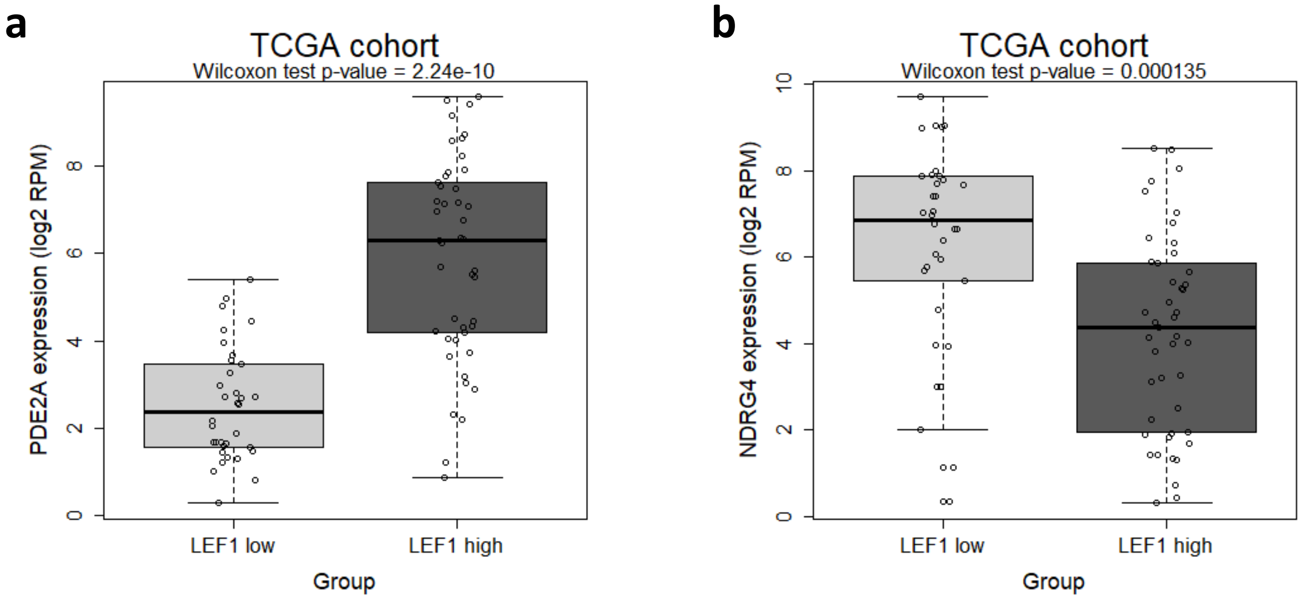


**Supplementary Figure S13. Expression levels of PDE2A** (a) **and NDRG4** (b) **in function of LEF1 expression level in the TCGA cohort.**

### Aberrant activation of Wnt/β-Catenin signaling pathway drives the expression of poor prognosis-associated microRNAs in adrenocortical cancer with a major impact on miR-139-5p and its host gene PDE2A

Justine Cristante[1], Soha Reda El Sayed[1], Josiane Denis[1], Ragazzon Bruno[2], Constanze Hantel[3,4], Olivier Chabre[1,5], Laurent Guyon[1#], Nadia Cherradi[1#]

#### Supplementary Tables

**Supplementary Table 1 :** qPCR TaqMan miRNA assay ID

| miRNA | TaqMan Assay ID |
|---|---|
| hsa-miR-139-5p | 002289 |
| hsa-miR-483-5p | 002338 |
| hsa-miR-483-3p | 002339 |
| hsa-miR-377-5p | 002128 |
| hsa-miR-377-3p | 000566 |
| hsa-miR-1185-2-3p | 476998_mat |
| hsa-miR-541-3p | 002201 |
| hsa-miR-320b | 002844 |
| hsa-miR-2110 | 121216_mat |
| hsa-miR-330-3p | 000544 |
| hsa-miR-513a-3p | 002091 |
| RNU48 | 001006 |

**Supplementary Table 2 :** qPCR primers

| Gene | Accession number | Forward primer | Reverse primer |
|---|---|---|---|
| HPRT | NM_000194.3 | 5'- ATGGACAGGACTGAACGTCTTGCT -3' | 5'- TTGAGCACACAGAGGGCTACAATG -3' |
| RPL13A | NM_012423.4 | 5'-TTAATTCCTCATGCGTTGCCTGCC-3' | 5'-TTCCTTGCTCCCAGCTTCCTATGT-3' |
| CTNNB1 (β-Cat) | NM_001904.4 | 5'-CTTCACCTGACAGATCCAAGTC-3' | 5'-CCTTCCATCCCTTCCTGTTTAG-3' |
| Axin2 | NM_004655.4 | 5'-CGAGTAGCCAAAGCGATCTAC-3' | 5'-GCTGCTTCTTGATGCCATCTC-3' |
| PDE2A | NM_002599.5 | 5'-GAGAGGCACCACTTTGCTCA-3' | 5'-ATGCGCTGATAGTCCTTCG-3' |
| NDRG4 | NM_020465.4 | 5'-ATGTGGGCCTCAACCACAAA-3' | 5'-GGAACTGGTACCCCTGAGGA-3' |

**Supplementary Table 3:** List of the 54 mature miRNAs whose expression was changed following β-Cat silencing. Selected miRNAs for RT-qPCR validation are indicated in bold.

| miR ID | MIMAT | Type of change |
|---|---|---|
| hsa-miR-504-5p | MIMAT0002875 | ↓ |
| **hsa-miR-330-3p** | MIMAT0000751 | ↓ |
| hsa-miR-182-3p | MIMAT0000260 | ↓ |
| hsa-miR-6772-3p | MIMAT0027445 | ↓ |
| hsa-miR-653-5p | MIMAT0003328 | ↓ |
| **hsa-miR-377-5p** | MIMAT0004689 | ↓ |
| hsa-miR-135a-3p | MIMAT0004595 | ↓ |
| hsa-miR-4443 | MIMAT0018961 | ↓ |
| hsa-miR-760 | MIMAT0004957 | ↓ |
| hsa-miR-769-3p | MIMAT0003887 | ↓ |
| hsa-miR-181c-5p | MIMAT0000258 | ↓ |
| hsa-miR-628-5p | MIMAT0004809 | ↓ |
| hsa-miR-653-3p | MIMAT0026625 | ↓ |
| hsa-miR-425-3p | MIMAT0001343 | ↓ |
| **hsa-miR-541-3p** | MIMAT0004920 | ↓ |
| hsa-miR-7977 | MIMAT0031180 | ↓ |
| hsa-miR-4326 | MIMAT0016888 | ↓ |
| **hsa-miR-320b** | MIMAT0005792 | ↓ |
| **hsa-miR-377-3p** | MIMAT0000730 | ↓ |
| hsa-miR-4753-3p | MIMAT0019891 | ↓ |
| hsa-miR-1306-5p | MIMAT0022726 | ↓ |
| **hsa-miR-139-5p** | MIMAT0000250 | ↓ |
| hsa-let-7i-3p | MIMAT0004585 | ↓ |
| hsa-miR-184 | MIMAT0000454 | ↓ |
| hsa-miR-1908-5p | MIMAT0007881 | ↓ |
| hsa-miR-92b-5p | MIMAT0004792 | ↓ |
| hsa-miR-135a-5p | MIMAT0000428 | ↓ |
| hsa-miR-532-5p | MIMAT0002888 | ↓ |
| hsa-miR-331-3p | MIMAT0000760 | ↓ |
| hsa-miR-363-3p | MIMAT0000707 | ↓ |
| **hsa-miR-513a-5p** | MIMAT0002877 | ↓ |
| hsa-miR-370-5p | MIMAT0026483 | ↓ |
| hsa-miR-589-5p | MIMAT0004799 | ↓ |
| hsa-miR-324-5p | MIMAT0000761 | ↓ |
| hsa-miR-125b-1-3p | MIMAT0004592 | ↓ |
| hsa-miR-489-3p | MIMAT0002805 | ↓ |
| **hsa-miR-1185-2-3p** | MIMAT0022713 | ↓ |
| hsa-miR-99a-3p | MIMAT0004511 | ↓ |
| hsa-miR-676-3p | MIMAT0018204 | ↓ |
| hsa-miR-191-3p | MIMAT0001618 | ↓ |
| **hsa-miR-2110** | MIMAT0010133 | ↓ |
| hsa-miR-1468-5p | MIMAT0006789 | ↓ |
| hsa-miR-26a-2-3p | MIMAT0004681 | ↑ |
| hsa-miR-450a-1-3p | MIMAT0022700 | ↑ |
| hsa-miR-1304-5p | MIMAT0005892 | ↑ |
| hsa-miR-3162-5p | MIMAT0015036 | ↑ |
| hsa-miR-561-3p | MIMAT0003225 | ↑ |
| hsa-miR-152-3p | MIMAT0000438 | ↑ |
| hsa-miR-145-3p | MIMAT0004601 | ↑ |
| hsa-let-7f-1-3p | MIMAT0004486 | ↑ |
| hsa-miR-431-3p | MIMAT0004757 | ↑ |
| hsa-miR-221-5p | MIMAT0004568 | ↑ |
| hsa-miR-202-5p | MIMAT0002810 | ↑ |
| hsa-miR-758-5p | MIMAT0022929 | ↑ |
